# A comprehensive study of light quality acclimation in *Synechocystis* sp. PCC 6803

**DOI:** 10.1101/2023.06.08.544187

**Authors:** Tomáš Zavřel, Anna Segečová, László Kovács, Martin Lukeš, Zoltán Novák, Anne-Christin Pohland, Milán Szabó, Boglárka Somogyi, Ondřej Prášil, Jan Červený, Gábor Bernát

**Affiliations:** Global Change Research Institute of the Czech Academy of Sciences, Brno, 603 00, Czechia; Institute of Plant Biology, HUN-REN Biological Research Centre, Szeged, 6726, Hungary; Centre Algatech, Institute of Microbiology of the Czech Academy of Sciences, Třeboň, 379 01, Czechia; HUN-REN Balaton Limnological Research Institute, Tihany, 8237, Hungary

**Keywords:** Cyanobacteria, Light harvesting, Light quality, Photomorphogenesis, Photosynthesis, State transitions

## Abstract

Cyanobacteria play a key role in primary production in both oceans and fresh waters and hold great potential for sustainable production of a large number of commodities. During their life, cyanobacteria cells need to acclimate to a multitude of challenges, including shifts in intensity and quality of incident light. Despite our increasing understanding of metabolic regulation under various light regimes, detailed insight into fitness advantages and limitations under shifting light quality has been missing. Here, we study photo-physiological acclimation in the cyanobacterium *Synechocystis* sp. PCC 6803 through the whole range of photosynthetically active radiation (PAR). Using LEDs with qualitatively different narrow spectra, we describe wavelength dependence of light capture, electron transport and energy transduction to main cellular pools. In addition, we describe processes fine-tuning light capture such as state transitions and efficiency of energy transfer from phycobilisomes to photosystems. We show that growth was the most limited under blue light due to inefficient light harvesting, and that many cellular processes are tightly linked to the redox state of the PQ pool, which was the most reduced under red light. The PSI-to-PSII ratio was low under blue photons, however, it was not the main growth-limiting factor, since it was even more reduced under violet and near far-red lights, where *Synechocystis* grew faster compared to blue light. Our results provide insight into the spectral dependence of phototrophic growth and can provide the foundation for future studies of molecular mechanisms underlying light acclimation in cyanobacteria, leading to light optimization in controlled cultivations.

## Introduction

Primary production both in natural aquatic systems and in laboratory- or industrial-scale cultivations largely depends on the ability of cells to acclimate to the surrounding environment. The factors affecting the fitness of phototrophic microorganisms include temperature, nutrient levels, pH, as well as both intensity and quality of the incident light. Among these, the impact of light quality, i.e. a specific characteristics of the underwater spectra, has been for long the least studied phenomenon. However, as evident from recent studies, the underwater spectrum is one of the most crucial factors that drive worldwide phytoplankton distribution (Grébert et al., 2018; Holtrop et al., 2021). Moreover, light quality has been shown to affect phototrophic production of commodities such as isoprene (Rodrigues et al., 2023).

Microalgae and cyanobacteria have developed numerous mechanisms to optimize light harvesting and energy transduction in conditions of either excessive or limited light availability. The short-term light acclimation includes state transitions (ST) (Calzadilla and Kirilovsky, 2020), decoupling of light-harvesting antenna (Tamary et al., 2012), non-photochemical quenching (NPQ) (Kirilovsky and Kerfeld, 2016), heat dissipation (Demmig-Adams and Adams, 2006), activation of futile cycles such as cycling of inorganic carbon between cells and the environment (Müller et al., 2019; Tchernov et al., 2003), scavenging of reactive oxygen species (Pospíšil, 2012), or shifts in transcriptome (Luimstra et al., 2020). The long-term acclimation involves complex changes in proteome (Jahn et al., 2018; Zavřel et al., 2019) including ratio of photosystem I (PSI) to photosystem II (PSII) (Luimstra et al., 2020), regulation of synthesis of energy-storing molecules such as glycogen (Cano et al., 2018) or lipids (Zavřel et al., 2018c) and modifications of photosynthetic antenna during chromatic acclimation (CA).

The latter process provides a great advantage in the spectrally-limited underwater environments; while many strains are restricted to certain spectral niches, the chromatic acclimators can efficiently harvest a broader spectrum of available light and, therefore, significantly increase the area and depth of suitable habitats (Sanfilippo et al., 2019). Up to date, eight types of CA have been recognized (CA0-7). CA1, present in *Synechocystis* sp. PCC 6803 (hereafter referred to as *Synechocystis*), is described below. During CA2, phycoerythrin (PE) is upregulated under green light. CA3 combines complementary upregulation of PE and phycocyanin (PC) under green and red lights, respectively. During CA4, the amounts of phycourobilin and phycoerythrobilin chromophores within phycobilisomes (PBS) are shifted under blue and green lights, without affecting PBS proteins. CA5 and CA6 are related to red/far-red light acclimation. During CA5 the light harvesting is secured by chlorophyll *d* incorporated in the thylakoid membrane instead of PC-containing PBS that are absent. During CA6 far-red light results in complex changes such as the induction of far-red shifted allophycocyanin and alternative photosystem proteins with a shift from chlorophyll *a* to *d*/*f* (Sanfilippo et al., 2019). CA0 and CA7 optimize yellow-green light absorption regulating rod-shaped PBS (CA0) and phycoerythrocyanin (CA7) (Hirose et al., 2019).

While the mechanisms underlying various types of CA are known to great detail, less attention has been paid to the implications of light quality shift on phytoplankton energetics and metabolism. The summary of metabolic changes in response to light signals is called photomorphogenesis (Bernát et al., 2021; Montgomery, 2016), or photoacclimation. Even though in cyanobacteria the response to individual wavelengths has been studied to some extent, only a few studies provided a complex understanding to cellular energetics under a light quality shift (Bernát et al., 2021). This is surprising, since the ability of phytoplankton to effectively harvest available light over a range of spectral niches ultimately determines its abundance in the environment (Holtrop et al., 2021).

This study provides a detailed report on light quality acclimation of *Synechocystis*, a type 1 chromatic acclimator. CA1 possess a specific i) green light-dependent redistribution of light energy between photosystems through the linker CpcL that binds PBS preferentially to PSI, and ii) a red light-dependent binding of the PC rods to the allophycocyanin (APC) core through a CpcG1 linker that helps to form canonical PBS (for more details see review (Sanfilippo et al., 2019)). The red/green cyanobacterial chromatic acclimation sensor (CcaS) that regulates also CpcL is common for type CA1-3 (Wang and Chen, 2022), whereas the CpcL linker is present in many strains that are even not classified as CA (Hirose et al., 2019). The light acclimation in *Synechocystis* thus partially reveals photoacclimation strategies of other phytoplankton species that do not possess CA. One example is a close *Synechocystis* relative *Cyanobium gracile* that adopted both common and unique photoacclimation traits (Bernát et al., 2021).

In general, the most favorable wavelengths for *Synechocystis* and many other cyanobacteria are those around the absorption maxima of the PBS (orange/red photons), whereas the least favorable are blue photons. The poor growth under blue light is a result of excitonic imbalance between PSII and PSI. The majority of blue photons is absorbed by PSI which is typically several times more abundant than PSII (Bernát et al., 2021; Luimstra et al., 2018; Murakami et al., 1997) and which binds more chlorophyll molecules in its structure (Netzer-El et al., 2019; Umena et al., 2011). The under-excitation of PSII limits oxygen evolution and linear electron transport rate (Luimstra et al., 2018) which further limits NADPH production, carbon fixation and ultimately cell division (Pfennig et al., 2023). On the other hand, orange-red light is harvested by PBS that transfer the excitation energy efficiently to PSII (Li et al., 2021) but less efficiently to PSI.

Despite the detailed understanding of blue- and red-light driven shifts in transcriptome or photosynthesis efficiency (Luimstra et al., 2020, 2018; Singh et al., 2010) far less is known about acclimation to other wavelengths. Recent works reported shifts in photosynthetic efficiency and pigmentation under near UV, green, and near far-red wavelengths (Bernát et al., 2021; Rodrigues et al., 2023). However, reports systematically comparing light acclimation across the whole visible light spectrum (Pfennig et al., 2023) are still scarce.

This study provides insight into spectral acclimation in a CA1 performing cyanobacterium, by addressing the regulation of light capture and energy transduction across the visible light spectrum (435-687 nm). The results show complex trends in the wavelength dependency of cell composition, energy transfer from PBS to PSII and PSI, state transitions, and non-photochemical quenching as well as in the redox state of the PQ pool. The results extend our understanding of phytoplankton limitation in natural environments and can shape the design of new strains as well as cultivation strategies in the controlled cultivation systems.

## Results

### Growth rate is tightly linked with the rate of electron flow

For the experiments, light intensity of nine narrow-band LEDs with peak wavelengths at 435-687 nm was set to 25 µmol photons m^−2^ s^−1^ (*Fig. 1a*). The photosynthetically usable radiation (PUR) as calculated from photosynthetically active radiation (PAR) and absorption spectra (*Fig. 1b*), was the highest under violet (435 nm) and blue light (465 nm, *Fig. 1c*). However, *Synechocystis* grew with the highest specific growth rate under orange/red LEDs (633-663 nm, *Fig. 1g*). These high growth rates were accompanied by high linear electron flow from PSII to PSI (LEF, *Fig. 1d*), cyclic electron flow around PSI (PSI-CEF, *Fig. 1e*), and respiratory electron flow (REF, *Fig. 1f*). Electron flow through PSI was limited at the donor side under blue, green, yellow and orange light (Y(ND) ∼ 0.6, *Fig. 1h*), while some minor acceptor side limitation (Y(NA) ∼ 0.07) occurred at 687 nm LEDs (*Fig. 1i*). No lag-phase was observed under any tested light (Supplementary Figure S1).

**Fig. 1.**
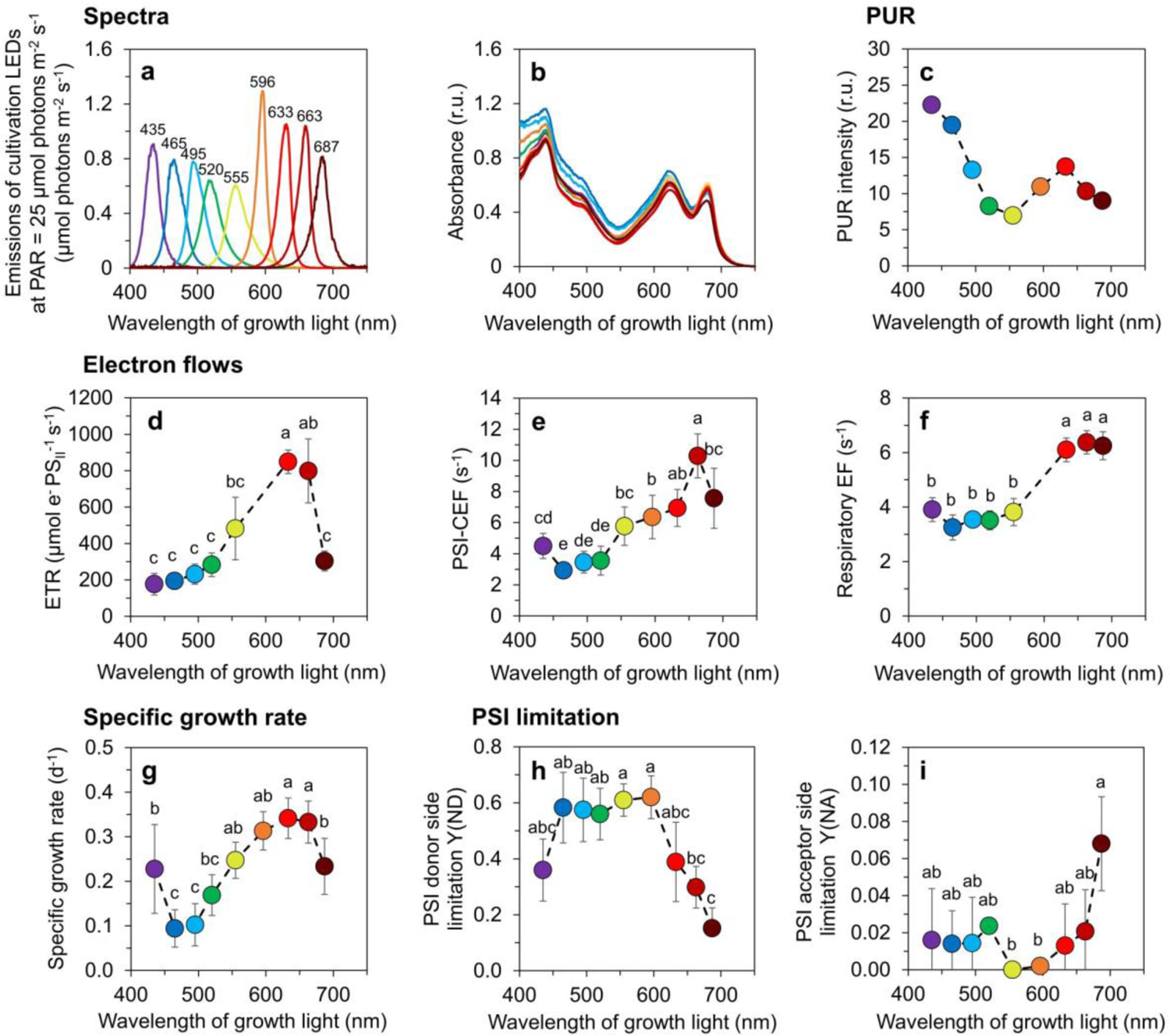
Growth rate and electron flows in *Synechocystis* cells grown under narrow-band cultivation lights. Emission spectra of narrow-band cultivation LEDs at photosynthetically active radiation (PAR) of 25 µmol photons m^−2^ s^−1^ (**a**), baseline corrected absorption spectra of *Synechocystis* cultures (**b**), photosynthetically usable radiation (PUR, **c**, see Eq. (6) for details), electron transport rate through PSII (ETR, **d**), cyclic electron flows around PSI (PSI-CEF, **e**), respiratory electron flow (**f**), specific growth rate (**g**), and donor side (Y(ND), **h**) and acceptor side (Y(NA), **i**) limitation of PSI in *Synechocystis* cells as cultivated under narrow-band LEDs. The values in panels **b** and **d**-**i** represent mean±SD (n = 3-6). Values in panel **b** (n=3) are shown without error bars for clarity. The letters above the symbols in panels **d-i** indicate statistically significant differences within each parameter (p<0.05).

### Increased electron flow under red light allows to accumulate more cellular reserves

The level of all pigments, including phycobilisomes (*Fig. 2a*), Chl *a* (*Fig. 2b*) and carotenoids (*Fig. 2c*) were relatively low under blue light (465 nm). In contrast, relatively high cellular content of PBS and carotenoids was observed under violet (435 nm), near far-red (687 nm) and also under yellow light (555 nm), where Chl *a* performed the highest level (*Fig. 2a-c*). The increased amount of carotenoids included echinenone, myxoxanthophyll, zeaxanthin, and synechoxanthin, whereas β-carotene was found to be less dependent of the cultivation wavelength (Supplementary Figure S2). The calculated carotenoids / Chl *a* ratio was the highest under violet and near far-red lights (Supplementary Figure S3). Despite dramatic changes in the content of light harvesting pigments, the functional absorption cross-section of PSII (σ_II_) was independent of the cultivation wavelength (Supplementary Figure S4).

**Fig. 2.**
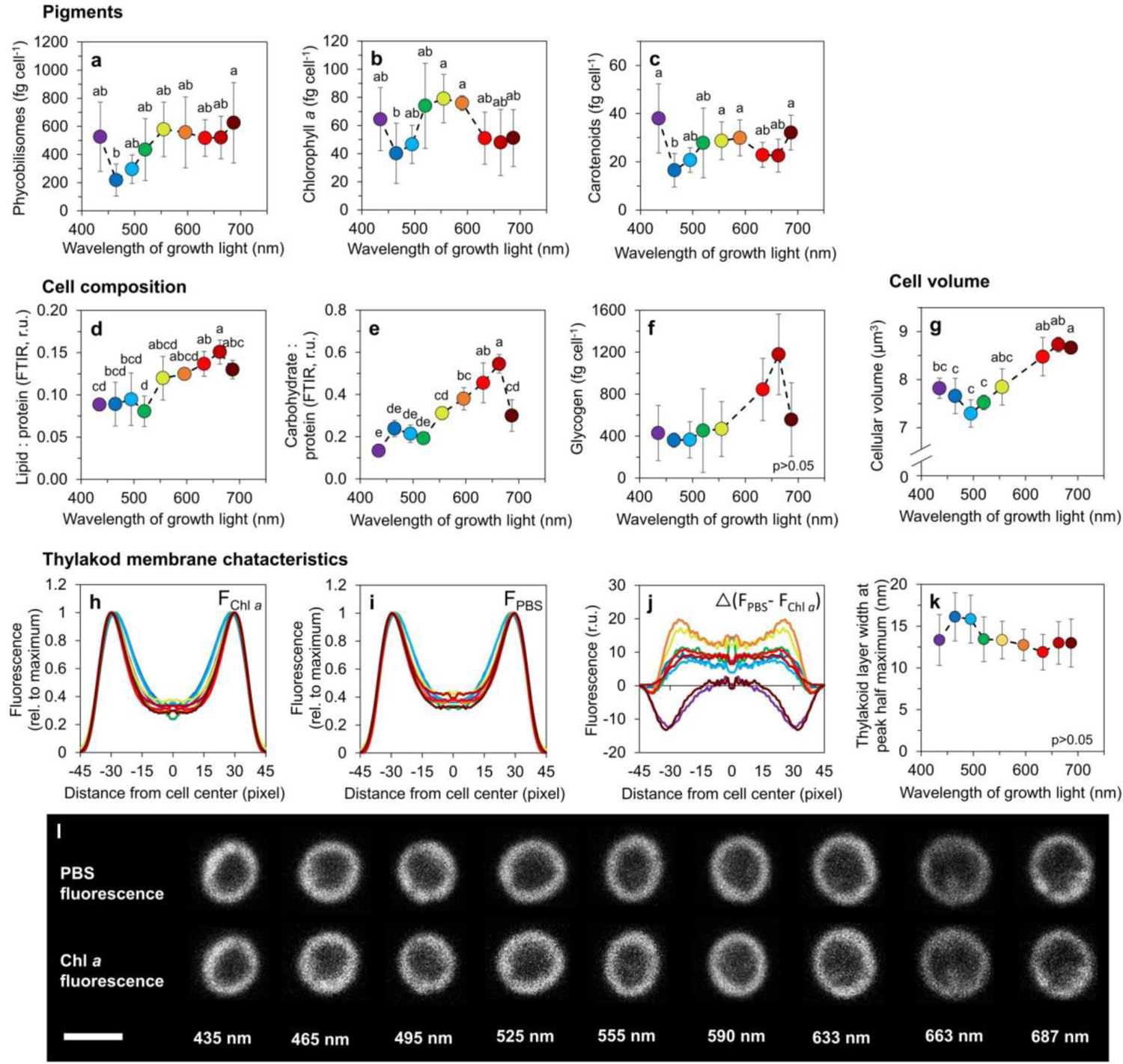
Macromolecular composition and morphology of *Synechocystis* cells grown under narrow-band cultivation lights. Content of phycobilisomes (**a**), chlorophyll *a* (***b***) and carotenoids (**c**) in *Synechocystis* cells, carbohydrate (**d**) and lipid (**e**) relative to protein, and cellular content of glycogen (**f**). Cell volume (**g**). Parameters derived from confocal microscopy imaging: autofluorescence profiles of chlorophyll *a* (F_Chl *a*_, **h**) and phycobilisomes (F_PBS_) **i**) across *Synechocystis* cells, difference between F_PBS_ and F_Chl *a*_ (**j**) and thickness of the thylakoid membrane, determined from F_Chl *a*_ as half maximum of fluorescence peaks (**k**). All values represent mean±SD (n = 3–5 (-**g**) / 58-113 (**h-k**)). Error bars in panels **h-k** are omitted for clarity. Confocal microscopy images of representative *Synechocystis* cells as cultivated under narrow-spectrum lights are shown in panel **I** (scale bar: 2 µm). The letters above the symbols in panels **a-g** and **k** indicate statistically significant differences within each parameter. (p<0.05).

Efficient light harvesting and, in turn, high electron flow under red light allowed *Synechocystis* to accumulate cellular reserves. The cellular pools of carbohydrates and lipids were upregulated under orange to red light (633-663 nm, *Fig. 2d-f*; Supplementary Figure S5). As a consequence, *Synechocystis* cells had the largest cell volume under these wavelengths. However, we note that high cellular volume was found also under 687 nm light, where the content of carbohydrates was reduced compared to 663 nm light (*Fig. 2g*).

### *Synechocystis* has limited options to fine-tune light harvesting under blue light

To further understand the fine-tuning of light harvesting, 3D fluorescence excitation-emission maps were recorded at 77K (*Fig 3*). Analysis of the fluorescence spectra allowed to determine the excitation energy transfer from PBS to PSII (PBS-PSII) and to PSI (PBS-PSI), and the PBS population that was functionally uncoupled from both PSII and PSI (PBS-free). In addition, the energy emitted by the Chl *a* antenna of PSII (Chl-PSII) and PSI (Chl-PSII) was determined; for details see *Fig. 3* and (eq. 7-12)). PBS-PSII was the highest under violet and near-far red light and was relatively low under blue, green and orange light (*Fig. 3a*). This pattern was the opposite to that of the donor-side limitation of PSI (*Fig. 1h*) and consistent with the wavelength dependence of Chl-PSII (Supplementary Figure S6). PBS-PSI was also low under blue to orange light and was the highest under 663 nm light, similarly to Chl-PSI (*Fig. 3b*, Supplementary Figure S6). PBS-free showed significant differences only at shorter wavelengths, i.e. between blue (495 nm, lowest) and violet lights (435 nm, highest, *Fig. 3c*). Analysis of the data revealed relatively low PSI / PSII ratios under both violet (435 nm) and near far-red lights (687 nm, *Fig. 3d*), as a consequence of both high Chl-PSII and low Chl-PSI under these wavelengths (Supplementary Figure S6).

**Fig. 3.**
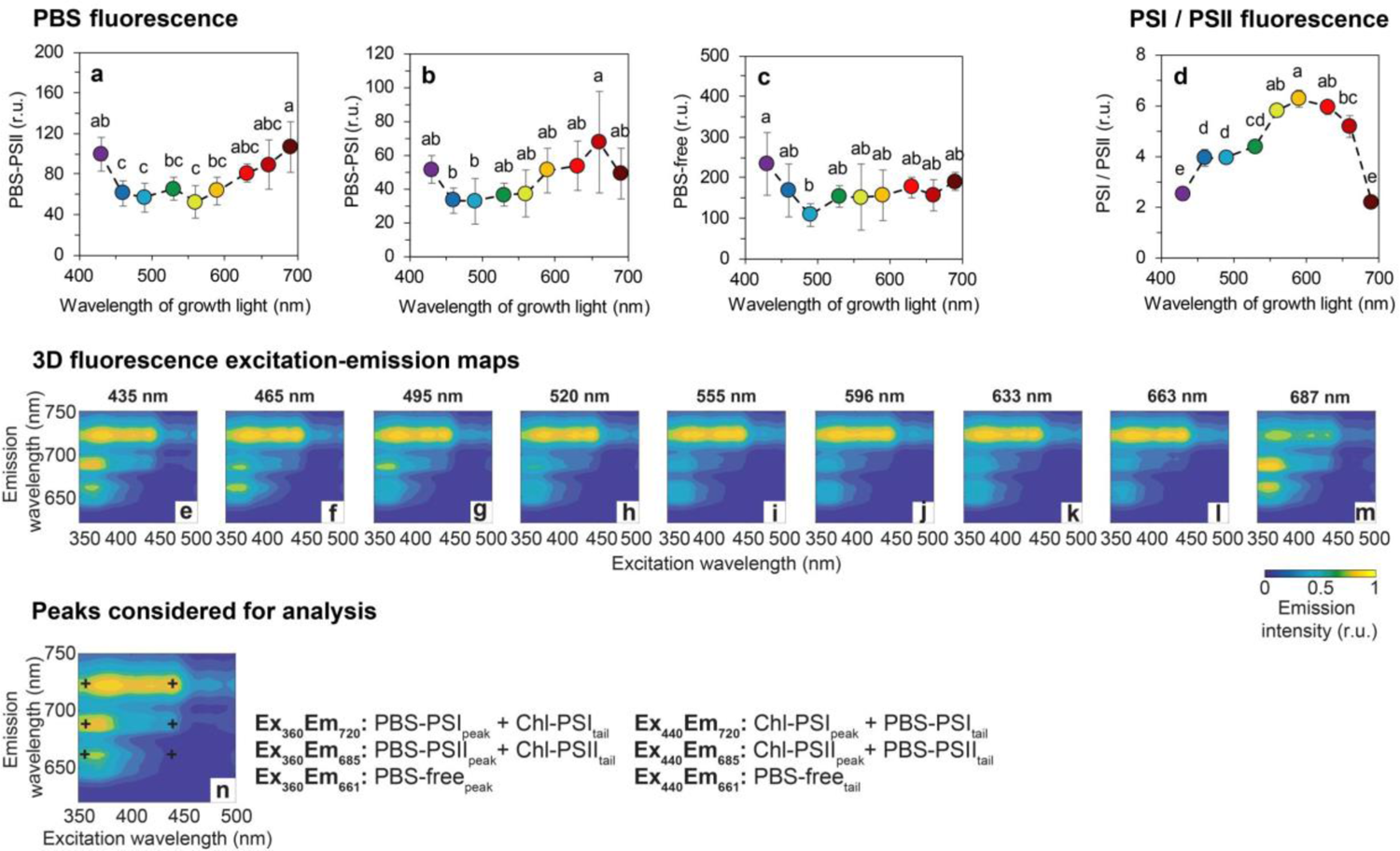
77K fluorescence excitation-emission maps and derived parameters under narrow-band cultivation lights. Fluorescence of phycobilisomes functionally attached to PSII (PBS-PSII, **a**), to PSI (PBS-PSI, **b**), and of phycobilisomes functionally uncoupled from both photosystems (PBS-free, **c**). PSI/PSII ratio (**d**); excitation-emission maps of *Synechocystis* cultures cultivated under narrow-band cultivation LEDs (**e-m**). The maps represent standardized averages (n = 3); error intervals are not shown for clarity. Representative fluorescence map with wavelengths used for analysis (**n**). For further details on the 77K spectra processing see Methods and Supplementary methods. The values in panels **a-d** represent mean±SD (n = 5-6). The letters above the symbols in panels **a-d** indicate statistically significant differences within each parameter.

Increased PSII abundance under violet and near far-red lights was further confirmed by the analysis of confocal micrographs that revealed higher autofluorescence of Chl *a* over PBS under these two particular cultivation wavelengths (*Fig. 2h-j*). Confocal microscopy also revealed the independence of the thickness of thylakoid membranes on the cultivation wavelength (*Fig. 2k*) and further confirmed PBS localization within the thylakoid membranes under all cultivation lights (*Fig. 2i,l*). This is in contrast to previous results in the cyanobacterium *Cyanobium gracile* where the PBSs were shown to detach from the thylakoid membrane layer under near far-red light (Bernát et al., 2021).

Plotting specific 2D fluorescence emission spectra upon 440 nm excitation (Supplementary Figures S7) revealed additional details, such as intensities of the 685 nm (F685) and 695 nm (F695) fluorescence emission bands, originating from CP43 and CP47 subunits of PSII, respectively (Remelli and Santabarbara, 2018), whose ratio can provide insight into PSII lifecycle and assembly state (Bečková et al., 2022). Although a slight increase in the F685/F695 ratio was found under blue and near far-red cultivation lights (Supplementary Figure S8), it was statistically not significant, suggesting similar PSII turnover rates under all tested wavelengths.

Besides studying the functional coupling of PBS to photosystems, the rates of state transitions (ST) were also determined, based on chlorophyll fluorescence transients. To induce *State 1* and *State 2*, *Synechocystis* cultures were illuminated by weak blue and red light, respectively (*Fig. 4c, f*). In general, *State 2 → State 1* transitions were always faster compared to *State 1 → State 2* transitions (*Fig. 4a-b*). Regarding the effect of cultivation wavelength, blue light (435-465 nm) was found to induce the slowest ST rates and, therefore, the least efficient capability to fine-tune light harvesting (*Fig. 4a-b*). In line with these results, NPQ as induced by strong blue actinic light (AL, *Fig. 4c, f*, eq. (1)) was also the weakest under blue cultivation light (*Fig. 4d-e*). Chlorophyll fluorescence kinetics under all cultivation wavelengths are shown in Supplementary Figure S9-S10.

**Fig. 4.**
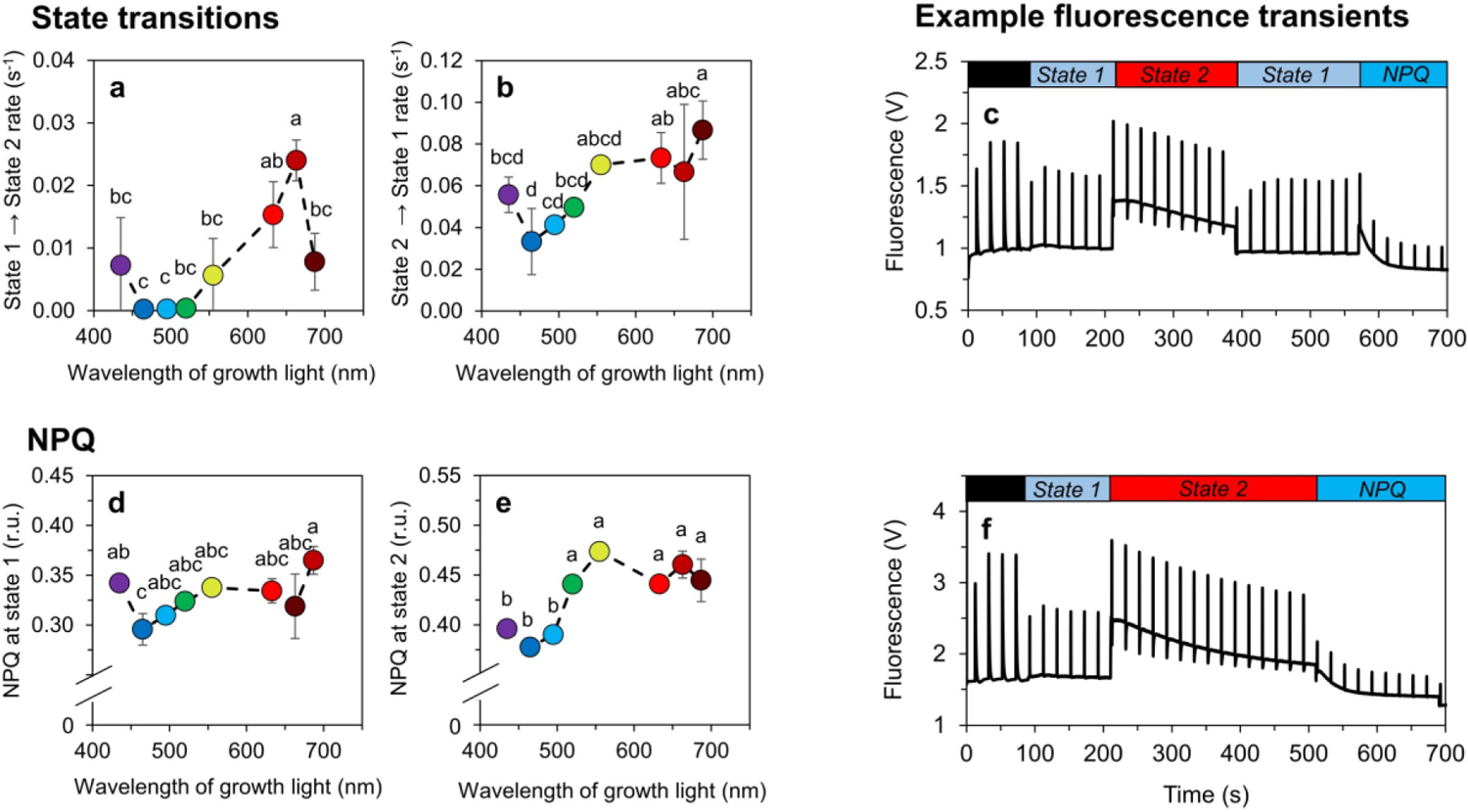
State transitions and NPQ in *Synechocystis* under narrow-band cultivation lights. Rates of *State 1 → State 2* (**a**) and *State 2 → State 1* (**b**) transitions induced by weak blue (480 nm, 80 µmol photons m^−2^ s^−1^) and red (625 nm, 50 µmol photons m^−2^ s^−1^) actinic light (AL), respectively. Non-photochemical quenching (NPQ) induced by strong blue AL (1 800 µmol photons m^−2^ s^−1^) in *State 1* (**d**) and *State 2* (**e**). Example PAM protocol used for the estimation of *State 2 → State 1* rate and NPQ in *State 1* is shown in panel **c**, example protocol for the estimation of *State 1 → State 2* rate and NPQ in *State 2* is shown in panel **f**. Prior to each measurement, *Synechocystis* cells were dark-acclimated for 15 min. Fluorescence recordings from all cultivation lights are summarized in Supplementary Figure S9-S10. The values in panels **a-b** and **d-e** represent mean±SD (n = 3), the letters above the symbols indicate statistically significant differences within each parameter (p<0.05).

### Red light leads to reduction of the PQ pool

Fast chlorophyll fluorescence transients (the so-called OJIP curves) can provide semi-quantitative information about the redox state of the PQ pool. In particular, an increase in the fluorescence yield at the J-level (F_J_, 2 ms), has been proven to be correlated with a more reduced PQ pool (Tóth et al., 2007). For quantitative analysis, the relative fluorescence yield at the J-level (V_J_, Eq. 2) has been used as a suitable parameter reflecting the redox state of the PQ pool (Tsimilli-Michael et al., 2009).

The calculated V_J_ values, as derived from OJIP curves recorded in both light- and dark-acclimated states (*Fig. 5a, c*) suggest a more reduced PQ pool under red light, and, on the contrary, more oxidized PQ pool under blue light (*Fig. 5b, d*), in accordance with previous works (Foyer et al., 2012). This result was further supported by reduction of parameter ψE_0_ and upregulation of parameter δR_0_ under red cultivation light (Supplementary Figure S11). These parameters reflect the efficiency of electron transport between Q_A_^-^ and PQ, and between plastoquinol (PQH_2_) and PSI, respectively (Stirbet et al., 2018).

**Fig. 5.**
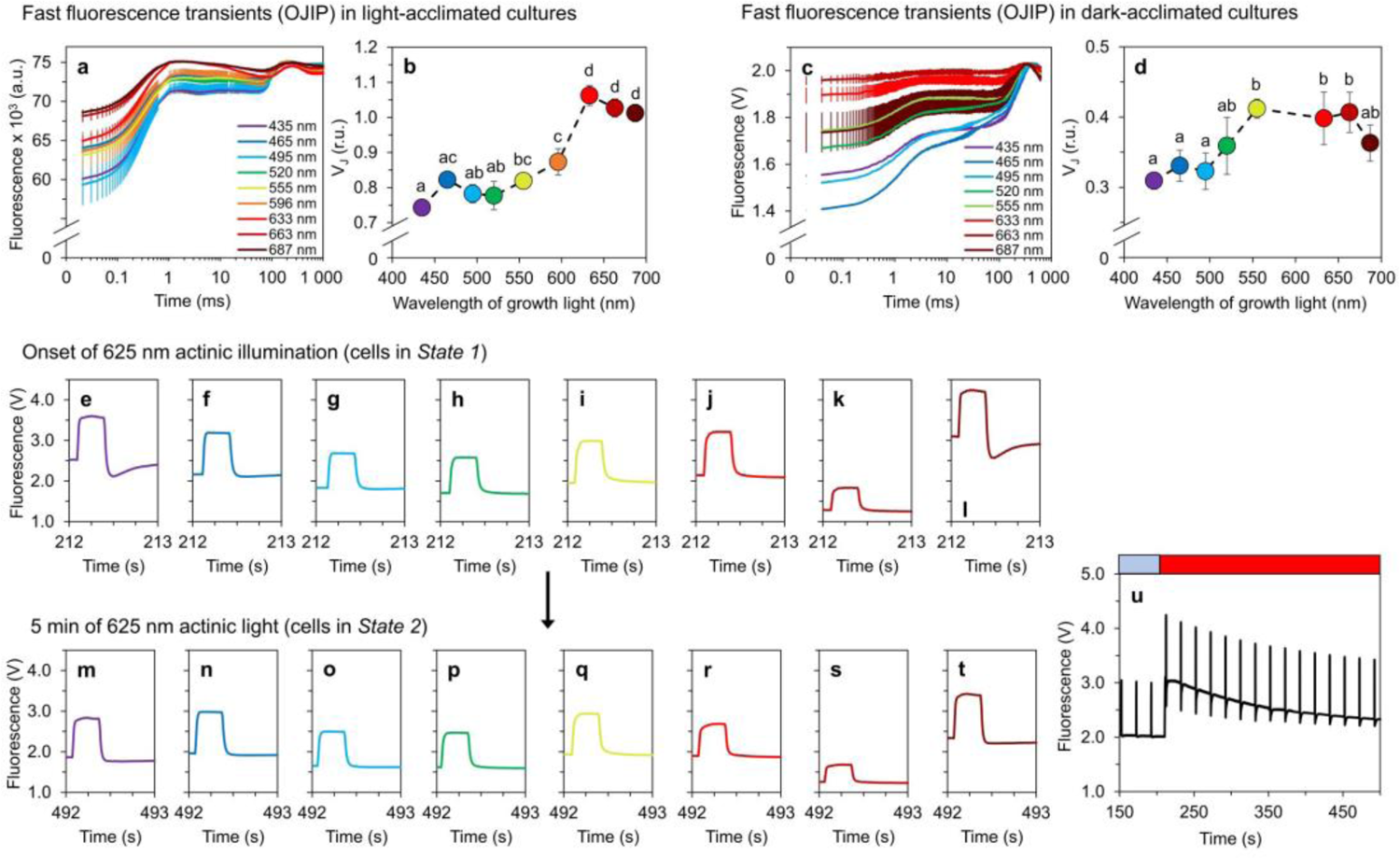
Redox state of the PQ pool and transient fluorescence drops after saturation pulses. Redox state of the PQ-pool was determined based on the relative fluorescence level at J point, V_J_ (**b**, **d**) derived from fast fluorescence kinetics (OJIP curves) in light-acclimated (**a**) and dark-acclimated *Synechocystis* cultures (**c**). Fluorescence traces during saturation pulses at the onset of 625 nm AL and after 5 min of 625 nm AL are summarized in panels **e-l** and **m-t**; respectively; an example pattern of a slow fluorescence kinetic is shown in panel **u**.

A closer analysis of the efficiency of linear electron transport revealed a temporal pattern in the PQ redox state. In cells cultivated under violet and near far-red light, the yield of the chlorophyll fluorescence dropped transiently below the steady-state level at the onset of red AL right after a saturation pulse (*Fig. 5e-l*). This drop was gradually decreased, and after 5 min of AL illumination it was negligible in all cultures (*Fig. 5m-t*). This suggests a temporal limitation of linear electron flow on the acceptor site (Tsuyama et al., 2004) under violet and near far-red lights. This was likely related to an increased relative PSII abundance under both abovementioned lights as well as to a reduced PSI abundance under violet light (Supplementary Figure S6). A summary of all changes in *Synechocystis*, as a response to individual monochromatic cultivation lights, is shown in *Fig. 6*.

**Fig. 6.**
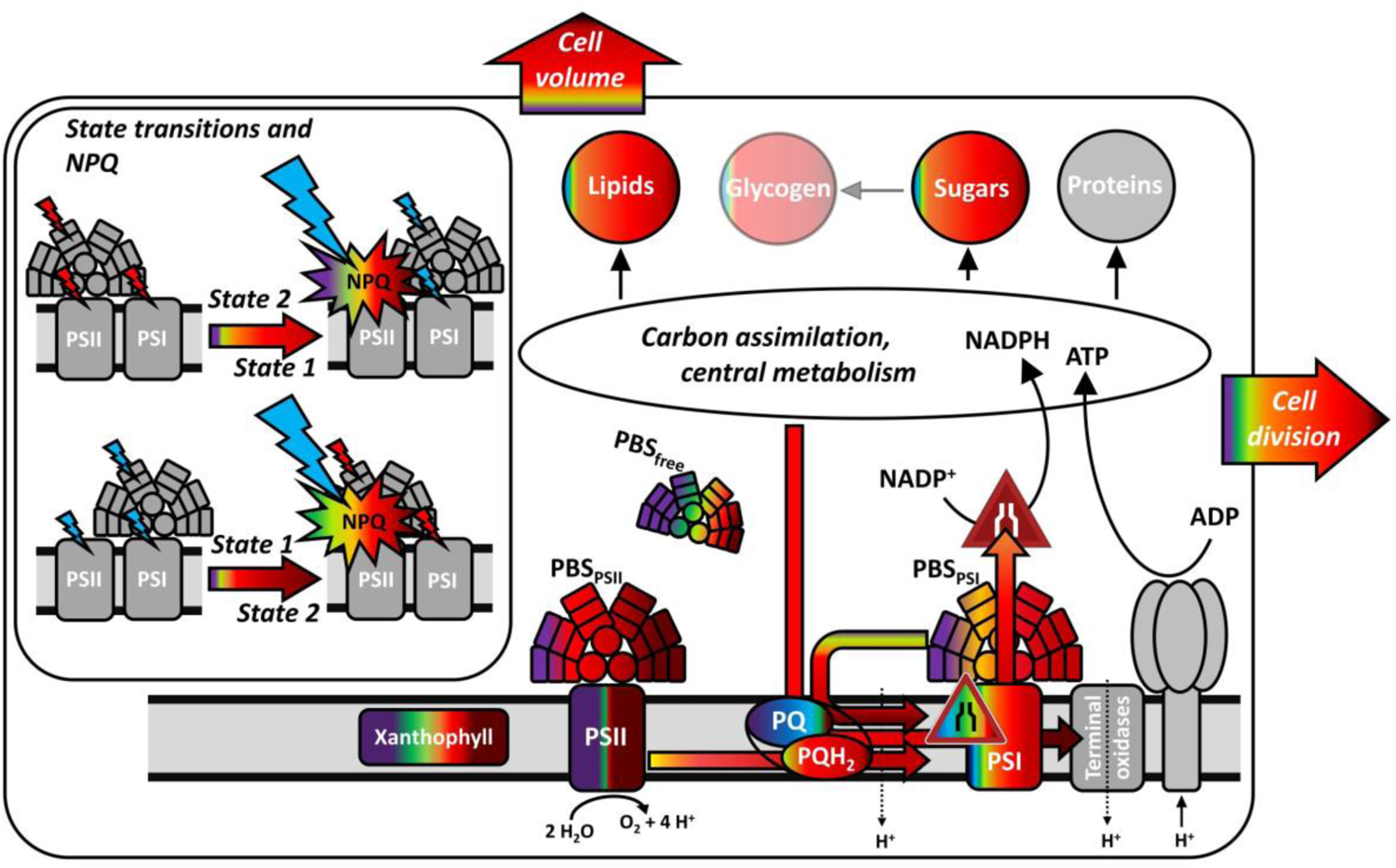
Summary of wavelength-dependent shifts in *Synechocystis* as found in this study. Individual processes and components are color-coded according to the narrow-band irradiance under which the corresponding processes/components were upregulated. Grey regions represent cellular components that were not determined (terminal oxidases, ATPase) or are depicted only for illustrative purpose (insert showing state transition). Components / processes dependent of cultivation wavelength are marked without transparency; glycogen is marked partially transparent as it was identified as independent of the wavelength of the cultivation light. Colored arrows represent the measured electron flows, black arrows represent pathways of which only the final sinks were determined.

All values represent means (n = 3), error bars in **a-d** represent standard deviations. Error bars in panels **e-t** are not shown for clarity. Fluorescence recordings from all cultivation lights are summarized in Supplementary Figures S9-S10. The letters above the symbols in panels **b** and **d** indicate statistically significant differences within each parameter (p<0.05).

## Discussion

In this work, the light quality acclimation in the model cyanobacterium *Synechocystis* was studied. The experimental setup covering the whole spectrum of visible light (*Fig. 1*) represents a significant improvement over many previous studies, where typically only two or three wavelengths were compared. Indeed, in case of PC-containing cyanobacteria such as *Synechocystis*, cultivation under blue and red light induces two fundamentally distinct light acclimation states which was also confirmed here (for details, see (Luimstra et al., 2020, 2018; Singh et al., 2009)). However, the current study shows that although some parameters shifted between blue and red cultivation lights almost linearly, the wavelength dependency of many other parameters was highly non-linear. In addition, the range of previously studied growth wavelengths was widened by addressing acclimations to violet and near far-red illumination. Both these cultivation lights were utilized better than blue light in *Synechocystis*, and both led to specific responses distinct from other wavelengths (such as reduced PSI/PSII ratio).

The relatively fast growth under near far-red light was presumably a result of the fact that even narrow-band LEDs emit light over a relatively broad range, in this particular case from about 625 nm (*Fig. 1*). Absorption of true far-red light (> 700 nm) is quite inefficient in *Synechocystis*, and growth under this light is as slow as under blue light (Wilde et al., 1997). Moreover, far-red light induces complex transcription shift that generally resembles stress response (Hübschmann et al., 2005) which is also common in blue light acclimation (Luimstra et al., 2020; Singh et al., 2009).

On the other hand, the increased growth rate under violet (435 nm) light compared to blue (465 nm) light was likely a result of higher PUR (*Fig. 1a-c*). Under violet light the higher growth rate was accompanied by slightly increased PSI-CEF (likely substituting a reduced PSI/PSII ratio, *Fig. 3d*) but not LEF, showing the importance of ATP generation for growth optimization. Both violet and near far-red light resulted in increased abundance of Chl-PSII and PBS-PSII (*Fig. 3*, Supplementary Figure S6) that, however, did not lead to higher LEF (*Fig. 1*). This can be related to reduced capabilities to fine-tune light harvesting (*Fig. 4*), again likely as a result of reduced PSI/PSII ratio. In addition, synthesis of xanthophylls but not β–carotene was upregulated under both violet and near far-red lights (Supplementary Figure S2) suggesting alterations in thylakoid membrane structure under both conditions (Rodrigues et al., 2023; Zakar et al., 2017).

Blue light is known to induce a system-wide acclimation response in *Synechocystis* (Luimstra et al., 2020, 2018; Singh et al., 2009). The growth limitation under blue light is a result of the inability to efficiently capture and transfer blue light to PSII, which leads to low LEF and insufficient production of reducing equivalents such as NADPH and/or Fd_red_ (Singh et al., 2009) as well as of ATP (Luimstra et al., 2020). The low LEF under blue photons was also found here (*Fig. 1*) and it was likely a key limiting factor for growth as well as for the accumulation of lipids and carbohydrates, which limitation eventually led to a decrease in cell volume (*Fig. 2*). Furthermore, under blue light we found strongly downregulated APC levels (Supplementary Figure S12), which further reduced efficiency of transfer of the captured light to both PSII and PSI (*Fig. 3)*. Transcript levels of APC genes were found upregulated under blue light in previous works, together with transcription of PBS degradation genes (Luimstra et al., 2020; Singh et al., 2009). It is therefore likely that the APC downregulation under blue light as reported here was related to post-transcriptional or (post-)translational regulation. The APC shortage can also explain the low kinetic rates of ST under blue and green cultivation light (*Fig. 4*). ST become relevant for light energy distribution only when PBS are assembled in its canonical form (i.e. CpcG1 form (Kondo et al., 2009)) which unlikely occurred under blue cultivation light that induced APC shortage (Supplementary Figure S12). Analogously, this APC downregulation can also explain reduced NPQ under blue light (*Fig. 4*).

The fact that the emission spectra of the 495 nm and 520 nm LEDs overlapped to a big extent, was a likely reason of similar values of many photosynthetic parameters recorded under these bluish lights, including growth rate. Many parameters changed dynamically above these growth wavelengths. Compared to the “495 nm” and “520 nm” light, we found a pronounced shift under green (555 nm) light in Chl *a* and carotenoid levels (*Fig. 2*), in particular in that of zeaxanthin, echinenone and synechoxanthin (Supplementary Figure S2). Since neither PSI nor PSII was upregulated under this green light (Supplementary Figure S6), the most likely explanation for the increased content of Chl *a* was its higher turnover rate, possibly compensating for the low absorbance of wavelengths around 550 nm (Fig. 1). Since free chlorophyll produces oxygen radicals (Krieger-Liszkay, 2004), such increased turnover rate expectedly triggers an overexpression of xanthophyll but not β–carotene, in line with our results (Supplementary Figure S2). We note that the increase xanthophyll levels under 687 nm cultivation light probably had a different origin, since Chl *a* level did not increase under that light (*Fig. 2*).

The increase in growth rate under 555 nm light, compared to the 495 nm and 520 nm light, can have multiple causes. First, due to its bandwidth, the emission spectrum of the 555 nm LEDs contained also some orange photons that are absorbed by PBS effectively and thus favor growth (*Fig. 1*). Second, the APC shortage present under blue light was not present under green light (Supplementary Figure S13). Third, the green light induces a formation of specific PBS form named CpcL-PBS which is able to bind to PSI through CpcL linker (Kondo et al., 2009). Shifting from blue to green light, the PBS were thus able to 1) absorb more light and 2) transfer the absorbed energy to both PSII and also to PSI *via* CpcL-PBS, resulting in higher LEF and PSI-CEF, as shown in *Fig. 1*.

The fastest growth of *Synechocystis* was observed under orange/red lights (peak wavelengths 633 nm and 663 nm), where LEF, PSI-CEF, and REF allowed to generate sufficient amount of ATP and reducing equivalents (*Fig. 1*). Our results show that this fast growth was achieved not only by high number of photosystems and light-harvesting antenna, or PBS coupling to PSII or PSI, but rather by a delicate interplay of many processes. These include, besides the abovementioned factors, also regulation of state transitions (*Fig. 4*) and synthesis of carbohydrates or lipids (*Fig. 2*). Indeed, the efficient electron flow through thylakoid membranes under red light led to increased accumulation of PQH_2_ over PQ, relative to other tested wavelengths (*Fig. 5*). The more reduced state of the PQ pool was likely related to increased ST (Calzadilla and Kirilovsky, 2020) as well as to the upregulation of genes of photosynthetic electron transport rate and other compartments (Foyer et al., 2012) as described in great detail previously (Luimstra et al., 2020; Singh et al., 2009; Wiltbank and Kehoe, 2019).

We note that our results are partly reliant on growth conditions, and the wavelength dependency of growth could change with shifts in temperature, salinity, and irradiance (Zavřel et al., 2017), or with the use of ammonia instead of nitrate as N source (Singh et al., 2009).

By addressing spectral dependency of light harvesting, electron transport and cellular energy storage, this work provides insight into spectral limitations of PC-rich cyanobacteria of CA1-type under controlled conditions. The results can navigate the design of new strains towards improved light utilization for the synthesis of targeted products. A call for such optimization has been announced (Klaus et al., 2022), and a proof-of-concept study addressing the effect of light quality on the synthesis of bulk chemicals has been provided recently (Rodrigues et al., 2023).

## Materials and Methods

### Inoculum cultures and experimental setup

All cultivations of *Synechocystis* sp. PCC 6803 GT-L (Zavřel et al., 2015a) were performed in a batch regime in 250 mL Erlenmeyer flasks on air at 24 °C in BG-11 medium (Rippka et al., 1979) under light-dark regime set to 14:10 h. All cultures were placed on a cultivation bench and were shaken daily to prevent excessive sedimentation of the cells. The inoculum cultures were cultivated under cool-white fluorescent lamps (25 µmol photons m^−2^ s^−1^). Prior to the cultivations under narrow-band LEDs, cultures were diluted with fresh BG-11 medium such that the optical density at 750 nm (Specord 210 plus, Analytik Jena, Germany) at the time of measurement was OD_750_ = 0.2 for all cultures.

During the growth experiments, illumination was secured by a home-built cultivation bench apparatus with 9 different types of narrow-band LEDs (for spectra, see *Fig. 1a*): FD-34UV-Y1 (peak wavelength: 435 nm), FD-3B-Y1 (465 nm), FD-32G-Y1 (495 nm), FD-3GY1 (520 nm), B08QCMC3K1 (555 nm), FD-3Y-Y1 (596 nm), FD-3R-Y1 (633 nm), FD-333R-Y1 (663 nm) and FD-34R-Y1 (687 nm). All LEDs but the 555 nm LED were manufactured by Shenzhen Fedy Technology Co. (Shenzhen, China). The 555 nm LED was manufactured by Nagulagu Co., Ltd. (Shenzhen, China). The PAR intensity of each illumination was set to 25 µmol photons m^−2^ s^−1^.

### Growth rate, cell size and cell composition

The specific growth rate was determined based on the OD_750_ values of exponentially growing cultures, using an exponential regression model. The cultures were cultivated until the late exponential growth phase (Supplementary Figure S1). Cell size was determined by an ImageStream MkII imaging flow cytometer (Amnis Corp., Seattle, WA, USA) using a previously described method (Zavřel et al., 2019). Briefly, samples were treated with 2% formaldehyde, incubated for 10 min at room temperature, and stored at –80°C. Prior to the analysis, samples were thawed at room temperature (∼ 30 min) and processed by flow cytometry to discriminate (i) focused objects and (ii) round objects (width/length ratio 0.9-1.0). During the cytometric analysis, pigment autofluorescence (excitation: 642 nm, detection: 642–745 nm) was also recorded to validate the selection of cells within all measured objects. Bright-field images were used for cell size analysis; cellular shape was assumed spherical.

The abundance of lipids and carbohydrates (relative to proteins) was estimated by Fourier-transformed infrared spectroscopy (FTIR), following a previously described protocol (Felcmanová et al., 2017). Briefly, 5 mL of the culture suspension was centrifuged (4 000 x *g*, 5 min), the supernatant was discarded, the pellet was freeze-dried, and dry pellet was analyzed by a Nicolet IS10 spectrometer (Thermo Fisher Scientific, Waltham, MA, USA).

The content of glycogen, Chl *a*, total carotenoids and PBS was determined by previously described protocols (Zavřel et al., 2018b, 2018a, 2015b). To distinguish the contribution of individual carotenoids to the total carotenoid pool, additional analysis was performed using high-performance liquid chromatography (HPLC) following a previously developed method (Bernát et al., 2021). Briefly, 10 mL cultures aliquots were harvested (Whatman glass microfiber filters GF/B; ⌀ 25 mm) and stored at – 80°C. Soluble pigments were extracted in 500 µL acetone and analyzed using a Shimadzu Prominence HPLC system (Shimadzu, Kyoto, Japan). Pigments separation was carried out using a Phenomenex Synergi 4 µm Hydro-RP 80 Å, LC Column 150 x 4.6 mm at 25°C. 20 µL aliquots were injected and the pigments were eluted by a linear gradient from solvent A (acetonitrile, water, trimethylamine; in a ratio of 9:1:0.01) to solvent B (ethyl acetate) at a flow rate of 1 mL min^-1^ (total time: 25 min). Pigments were identified according to the respective retention times and absorption spectra, and quantified by integrated chromatographic peak areas.

### Photosynthetic Activity Measurements

PSII activity was probed by MULTI-COLOR-PAM (MC-PAM; Walz, Effeltrich, Germany) using both slow (*SP-Analysis*) and fast fluorescence induction kinetics (*Fast Acquisition*), 625 nm measuring light (ML) and 15 min of dark acclimation.

During slow kinetics, actinic light (AL) of 480 nm (inducing *State 1*) or 625 nm (inducing *State 2* (Calzadilla and Kirilovsky, 2020)) was used, with intensities of 80 µmol photons m^−2^ s^−1^ and 50 µmol photons m^−2^ s^−1^, respectively (both were sufficiently high to induce fluorescence transients). To induce NPQ, 480 nm light with an intensity 1 800 µmol photons m^−2^ s^−1^ was additionally used. NPQ was calculated from maximal fluorescence value in dark (F_max_) and at the end of the high light period (F_max’_):

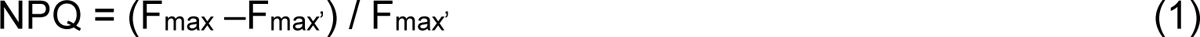

To calculate ETR, the functional absorption cross-section of PSII (σ_II_) was determined, using the default MC-PAM script *Sigma1000cyano* and a fitting protocol. In MC-PAM, the wavelength of AL defines also wavelength of saturation pulses (SP). It has to be noted that even the highest SP intensity (SP-int=20) at 480 nm was not strong enough to provide full PSII saturation, in contrast to SP of 625 nm (Supplementary Figure S13).

Fast fluorescence transients (OJIP curves) were recorded in both dark-acclimated and light-acclimated states, using MC-PAM and AquaPen (Photon System Instruments, Czechia) fluorometers and 625 nm / 620 nm saturation pulses, respectively. The redox state of PQ pool was estimated qualitatively based on the relative fluorescence yield at J point, V_J_ (Tóth et al., 2007; Tsimilli-Michael et al., 2009):

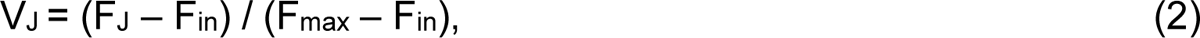

where F_J_ is fluorescence at the J-point of the OJIP curve (2 ms), and F_in_ and F_max_ are the initial (determined at time zero) and maximal fluorescence yield, respectively. The efficiency of e^-^ transport from Q_A_^-^ to PQ (ψE_0_) and from PQH_2_ to final acceptors (δR_0_) was calculated from V_J_ and V_I_ (relative fluorescence at I-level of the OJIP curve at 30 ms) according to (Stirbet et al., 2018):

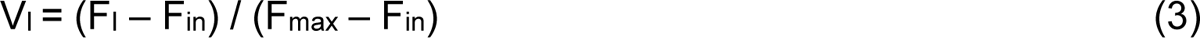

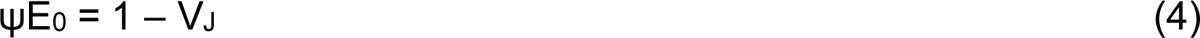

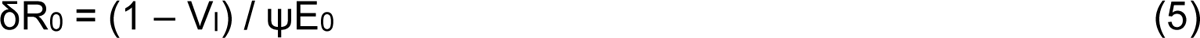

PSI kinetics was probed by the Dual-PAM-100 fluorometer (Walz, Effeltrich, Germany). Culture aliquots were filtered through glass fiber filters (GF/B, Whatman). Wet filters were placed between two microscope glass slides embedded in a DUAL-B leaf holder (Supplementary Figure S14) and dark-acclimated for 10 min. PSI limitation on both donor and acceptor sides was probed by *SP-analysis* mode (using *Fluo+P700 Measuring mode*). The rate of electrons transport through PSI was measured by *FastAcquisition* mode in the absence of inhibitors to estimate total electron flow through PSI, in the presence of 10 µM DCMU to estimate PSI-CEF (by inhibiting electron flow through PSII) and in the presence of 10 µM DCMU + 100 µM methyl-viologen (MV, a high affinity acceptor of PSI electrons) to quantify respiratory electron flow, by inhibiting PSII by DCMU and effectively preventing PSI-CEF by MV.

### Absorption and excitation-emission spectroscopy

Whole cell absorption spectra were recorded using a Specord 210 Plus spectrophotometer (Analytik Jena, Jena, Germany). To correct for light scattering by the cellular matter, four slices of tracing paper were placed in front of both sample and reference cuvettes. The recorded (offset-corrected) spectra were used to calculate photosynthetically usable radiation (PUR) for each narrow-band LED:

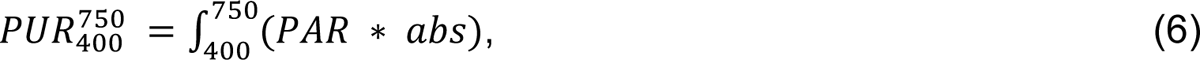

Where *PAR* and *abs* represent photosynthetically active radiation spectra of the cultivation LEDs and absorbance spectra of the corresponding *Synechocystis* cultures, respectively, in the range of 400-750 nm.

Fluorescence excitation-emission maps were recorded using a Jasco FP-8550 spectrofluorometer (Jasco, Tokyo, Japan) at 77K. 10 mL culture aliquots were filtered (GF/B filters, Whatman, Maidstone, UK), flash-frozen in liquid nitrogen and stored at −80 °C. Right before the measurement, a slice from each filter (∼1 cm × 0.3 cm) was cut to fit the metal holder of the transparent finger of the Dewar flask. 3D spectra maps were recorded over the excitation and emission range 350 nm - 650 nm (step: 5 nm) and 620 nm - 800 nm (step: 0.5 nm, bandwidth 5 nm, scan speed 1 000 nm min^-1^, sensitivity low), respectively. To distinguish fluorescence originating in PBS-PSI, PBS-PSI, PBS-free as well as chl *a* fluorescence originating either in PSII (Chl-PSII) or PSI (Chl-PSI), the following equations were used (Luimstra et al., 2020; Zavřel et al., 2021):

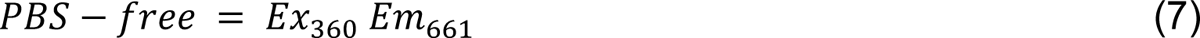

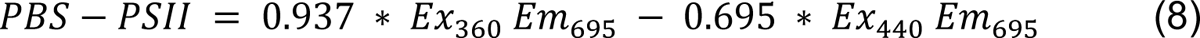

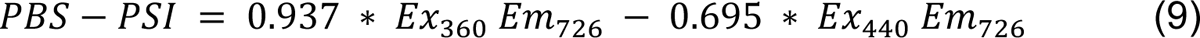

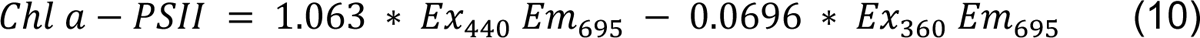

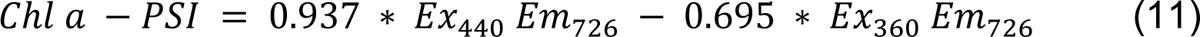

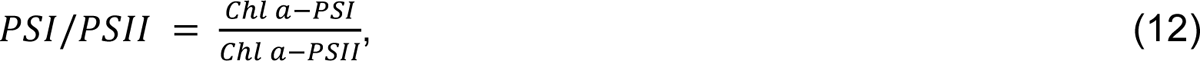

where Ex_360_ and Ex_440_ represent excitation at 360 nm and 440 nm (preferentially exciting PBS and Chl *a*, respectively), and Em_661_, Em_695_ and Em_726_ represent fluorescence emission by PBS, PSII and PSI, respectively as considered for the analysis. The coefficients in Eq. (7-12) provide correction for fluorescence emission tails, leaking between 360 and 440 nm (PBS and Chl *a* fluorescence tail, respectively). The coefficients were determined based on a comparison of fluorescence maps of *Synechocystis* 6803 WT and PAL mutant lacking PBS (Ajlani and Vernotte, 1998) (Supplementary Figure S15). Further details of the selection of particular excitation-emission wavelengths are provided in *Fig. 3*.

### Confocal microscopy

Microscopy imaging was performed using a TCS SP8 DMI confocal laser scanning microscope (Leica Microsystems Inc. Wetzlar, Germany) equipped with a HC PL APO CS2 63×/1.4 oil immersion objective and a TD 488/552/638 main beam splitter. 10 mL culture aliquots were centrifuged (2500 x *g*, 5 min, 25 °C), the supernatant was partially discarded and 10 µL of the concentrated sample was immobilized on a thin (<1 mm) layer of solid BG-11 agar and placed upside down on a cover slide. Chl *a* and PBS autofluorescence were excited using 488 nm and 638 nm lasers, respectively, and detected over 690–790 nm and 650–680 nm spectral windows. The recorded images were processed by using a self-developed Matlab script that allowed for evaluation of both Chl *a* and PBS fluorescence profiles within each identified cell (using 5° step, each cell was divided into 72 circular sectors around its geometrical center).

### Statistical analysis

To identify parameters that varied significantly among the applied light conditions, statistical methods using R Statistical Software (R Core Team, 2022) were applied as follows: ANOVA with Tukey HSD post-hoc test (Felipe de Mendiburu and Muhammad Yaseen, 2020) for cases when the homogeneity of variance (Fox and Weisberg, 2019) and normality of the data set was met; Kruskal-Wallis test followed by Multiple comparisons (Giraudoux et al., 2023) for cases when only the homogeneity of datás variance was met; Welch one-way test followed by Pairwise t-tests with BH correction for cases when only the normality of the data was met. The number of replicates was 3-9 for all growth lights throughout all experiments. The p-value was set to 0.05.

## Data Availability

The data underlying this article are available in the Figshare repository at 10.6084/m9.figshare.23330363.v1.

## Acknowledgement

We thank Tímea Szabó, Balázs Németh, and Máté Burányi for their technical assistance.

## Author Contributions

T.Z., and G.B. designed and performed experiments and/or generated material; T.Z., A.S., L.K., M.L., Z.N., A-C.P., M.S., B.S., O.P., J.Č., and G.B. designed experiments, analyzed data, and discussed results; T.Z. wrote the initial draft of the article; A.S., L.K., M.L., Z.N., A-C.P., M.S., B.S., O.P., J.Č., and G.B. contributed to finalizing the article.

## References

Ajlani, G., Vernotte, C., 1998. Construction and characterization of a phycobiliprotein-less mutant of Synechocystis sp. PCC 6803. Plant Mol. Biol. 37, 577–580.

Bečková, M., Sobotka, R., Komenda, J., 2022. Photosystem II antenna modules CP43 and CP47 do not form a stable ‘no reaction centre complex’ in the cyanobacterium Synechocystis sp. PCC 6803. Photosynth. Res. 152, 363–371.

Bernát, G., Zavřel, T., Kotabová, E., Kovács, L., Steinbach, G., Vörös, L., Prášil, O., Somogyi, B., Tóth, V.R., 2021. Photomorphogenesis in the Picocyanobacterium Cyanobium gracile Includes Increased Phycobilisome Abundance Under Blue Light, Phycobilisome Decoupling Under Near Far-Red Light, and Wavelength-Specific Photoprotective Strategies. Front. Plant Sci. 12, 1–16.

Calzadilla, P.I., Kirilovsky, D., 2020. Revisiting cyanobacterial state transitions. Photochem. Photobiol. Sci. 19, 585–603.

Cano, M., Holland, S.C., Artier, J., Burnap, R.L., Ghirardi, M., Morgan, J.A., Yu, J., 2018. Glycogen Synthesis and Metabolite Overflow Contribute to Energy Balancing in Cyanobacteria. Cell Rep. 23, 667–672.

Demmig-Adams, B., Adams, W.W., 2006. Photoprotection in an ecological context: the remarkable complexity of thermal energy dissipation. New Phytol. 172, 11– 21.

Felcmanová, K., Lukeš, M., Kotabová, E., Lawrenz, E., Halsey, K.H., Prášil, O., 2017. Carbon use efficiencies and allocation strategies in Prochlorococcus marinus strain PCC 9511 during nitrogen-limited growth. Photosynth. Res. 134, 71–82.

Felipe de Mendiburu, Muhammad Yaseen, 2020. agricolae: Statistical Procedures for Agricultural Research.

Fox, J., Weisberg, S., 2019. An R Companion to Applied Regression, Third. ed. Sage, Thousand Oaks {CA}.

Foyer, C.H., Neukermans, J., Queval, G., Noctor, G., Harbinson, J., 2012. Photosynthetic control of electron transport and the regulation of gene expression. J. Exp. Bot. 63, 1637–1661.

Giraudoux, P., Antonietti, J.-P., Beale, C., Groemping, U., Lancelot, R., Pleydell, D., Treglia, M., 2023.pgirmess: Spatial Analysis and Data Mining for Field Ecologists.

Grébert, T., Doré, H., Partensky, F., Farrant, G.K., Boss, E.S., Picheral, M., Guidi, L., Pesant, S., Scanlan, D.J., Wincker, P., Acinas, S.G., Kehoe, D.M., Garczarek, L., 2018. Light color acclimation is a key process in the global ocean distribution of Synechococcus cyanobacteria. Proc. Natl. Acad. Sci. U. S. A. 115, E2010– E2019.

Hirose, Y., Chihong, S., Watanabe, M., Yonekawa, C., Murata, K., Ikeuchi, M., Eki, T., 2019. Diverse Chromatic Acclimation Processes Regulating Phycoerythrocyanin and Rod-Shaped Phycobilisome in Cyanobacteria. Mol. Plant 12, 715–725.

Holtrop, T., Huisman, J., Stomp, M., Biersteker, L., Aerts, J., Grébert, T., Partensky, F., Garczarek, L., van der Woerd, H.J., 2021. Vibrational modes of water predict spectral niches for photosynthesis in lakes and oceans. Nat. Ecol. Evol. 5, 55–66.

Hübschmann, T., Yamamoto, H., Gieler, T., Murata, N., Börner, T., 2005. Red and far-red light alter the transcript profile in the cyanobacterium Synechocystis sp. PCC 6803: impact of cyanobacterial phytochromes. FEBS Lett. 579, 1613–8.

Jahn, M., Vialas, V., Karlsen, J., Ka, L., Uhle, M., Hudson, E.P., 2018. Growth of Cyanobacteria Is Constrained by the Abundance of Light and Carbon Assimilation Proteins. Cell Rep. 25, 478–486.

Kirilovsky, D., Kerfeld, C.A., 2016. Cyanobacterial photoprotection by the orange carotenoid protein. Nat. Plants 2.

Klaus, O., Hilgers, F., Nakielski, A., Hasenklever, D., Jaeger, K.E., Axmann, I.M., Drepper, T., 2022. Engineering phototrophic bacteria for the production of terpenoids. Curr. Opin. Biotechnol. 77, 102764.

Kondo, K., Mullineaux, C.W., Ikeuchi, M., 2009. Distinct roles of CpcG1-phycobilisome and CpcG2-phycobilisome in state transitions in a cyanobacterium Synechocystis sp. PCC 6803. Photosynth. Res. 99, 217–225.

Krieger-Liszkay, A., 2004. Singlet oxygen production in photosynthesis. J. Exp. Bot. 56, 337–346.

Li, M., Ma, J., Li, X., Sui, S.-F., 2021. In situ cryo-ET structure of phycobilisome–photosystem II supercomplex from red alga. Elife 10.

Luimstra, V.M., Schuurmans, J.M., Hellingwerf, K.J., Matthijs, H.C.P., Huisman, J., 2020. Blue light induces major changes in the gene expression profile of the cyanobacterium Synechocystis sp. PCC 6803. Physiol. Plant. 170, 10–26.

Luimstra, V.M., Schuurmans, J.M., Verschoor, A.M., Hellingwerf, K.J., Huisman, J., Matthijs, H.C.P., 2018. Blue light reduces photosynthetic efficiency of cyanobacteria through an imbalance between photosystems I and II. Photosynth. Res. 138, 177–189.

Montgomery, B.L., 2016. Mechanisms and fitness implications of photomorphogenesis during chromatic acclimation in cyanobacteria. J. Exp. Bot. 67, 4079–4090.

Müller, S., Zavřel, T., Červený, J., 2019. Towards a quantitative assessment of inorganic carbon cycling in photosynthetic microorganisms. Eng. Life Sci. 19, 955–967.

Murakami, A., Kim, S.J., Fujita, Y., 1997. Changes in Photosystem Stoichiometry in Response to Environmental Conditions for Cell Growth Observed with the Cyanophyte Synechocystis PCC 6714. Plant Cell Physiol. 38, 392–397.

Netzer-El, S.Y., Caspy, I., Nelson, N., 2019. Crystal Structure of Photosystem I Monomer From Synechocystis PCC 6803. Front. Plant Sci. 9.

Pfennig, T., Kullmann, E., Zavřel, T., Nakielski, A., Ebenhöh, O., Červený, J., Bernát, G., Matuszyńska, A., 2023. Shedding Light On Blue-Green Photosynthesis: A Wavelength-Dependent Mathematical Model Of Photosynthesis In Synechocystis sp. PCC 6803. bioRxiv.

Pospíšil, P., 2012. Molecular mechanisms of production and scavenging of reactive oxygen species by photosystem II. Biochim. Biophys. Acta - Bioenerg. 1817, 218– 231.

R Core Team, 2022.R: A Language and Environment for Statistical Computing.

Remelli, W., Santabarbara, S., 2018. Excitation and emission wavelength dependence of fluorescence spectra in whole cells of the cyanobacterium Synechocystis sp. PPC6803: Influence on the estimation of Photosystem II maximal quantum efficiency. BBA - Bioenerg. 1859, 1207–1222.

Rippka, R., Deruelles, J., Waterbury, J.B., Herdman, M., Stanier, R.Y., 1979. Generic Assignments, Strain Histories and Properties of Pure Cultures of Cyanobacteria. Microbiology 111, 1–61.

Rodrigues, J.S., Kovács, L., Lukeš, M., Höper, R., Steuer, R., Červený, J., Lindberg, P., Zavřel, T., 2023. Characterizing isoprene production in cyanobacteria – Insights into the effects of light, temperature, and isoprene on Synechocystis sp. PCC 6803. Bioresour. Technol. 380, 129068.

Sanfilippo, J.E., Garczarek, L., Partensky, F., Kehoe, D.M., 2019. Chromatic acclimation in cyanobacteria: A diverse and widespread process for optimizing photosynthesis. Annu. Rev. Microbiol. 73, 407–433.

Singh, A.K., Bhattacharyya-Pakrasi, M., Elvitigala, T., Ghosh, B., Aurora, R., Pakrasi, H.B., 2009. A systems-level analysis of the effects of light quality on the metabolism of a cyanobacterium. Plant Physiol. 151, 1596–608.

Singh, S.P., Der, D.H.Ã., Sinha, R.P., 2010. Cyanobacteria and ultraviolet radiation (UVR) stress: Mitigation strategies. Ageing Res. Rev. 9, 79–90.

Stirbet, A., Lazár, D., Kromdijk, J., Govindjee, 2018. Chlorophyll a fluorescence induction: Can just a one-second measurement be used to quantify abiotic stress responses? Photosynthetica 56, 86–104.

Tamary, E., Kiss, V., Nevo, R., Adam, Z., Bernát, G., Rexroth, S., Rögner, M., Reich, Z., 2012. Structural and functional alterations of cyanobacterial phycobilisomes induced by high-light stress. Biochim. Biophys. Acta - Bioenerg. 1817, 319–327.

Tchernov, D., Silverman, J., Luz, B., Reinhold, L., Kaplan, A., 2003. Massive light-dependent cycling of inorganic carbon between oxygenic photosynthetic microorganisms and their surroundings. Photosynth. Res. 77, 95–103.

Tóth, S.Z., Schansker, G., Strasser, R.J., 2007. A non-invasive assay of the plastoquinone pool redox state based on the OJIP-transient. Photosynth. Res. 93, 193–203.

Tsimilli-Michael, M., Stamatakis, K., Papageorgiou, G.C., 2009. Dark-to-light transition in Synechococcus sp. PCC 7942 cells studied by fluorescence kinetics assesses plastoquinone redox poise in the dark and photosystem II fluorescence component and dynamics during state 2 to state 1 transition. Photosynth. Res. 99, 243–255.

Tsuyama, M., Shibata, M., Kawazu, T., Kobayashi, Y., 2004. An Analysis of the Mechanism of the Low-wave Phenomenon of Chlorophyll Fluorescence. Photosynth. Res. 81, 67–76.

Umena, Y., Kawakami, K., Shen, J.-R., Kamiya, N., 2011. Crystal structure of oxygen-evolving photosystem II at a resolution of 1.9 Å. Nature 473, 55–60.

Wang, F., Chen, M., 2022. Chromatic Acclimation Processes and Their Relationships with Phycobiliprotein Complexes. Microorganisms 10, 1562.

Wilde, A., Churin, Y., Schubert, H., Börner, T., 1997. Disruption of a Synechocystis sp. PCC 6803 gene with partial similarity to phytochrome genes alters growth under changing light qualities. FEBS Lett. 406, 89–92.

Wiltbank, L.B., Kehoe, D.M., 2019. Diverse light responses of cyanobacteria mediated by phytochrome superfamily photoreceptors. Nat. Rev. Microbiol. 17, 37–50.

Zakar, T., Herman, E., Vajravel, S., Kovacs, L., Knoppová, J., Komenda, J., Domonkos, I., Kis, M., Gombos, Z., Laczko-Dobos, H., 2017. Lipid and carotenoid cooperation-driven adaptation to light and temperature stress in Synechocystis sp. PCC6803. Biochim. Biophys. Acta - Bioenerg. 1858, 337–350.

Zavřel, T., Chmelík, D., Sinetova, M.A., Červený, J., 2018a. Spectrophotometric Determination of Phycobiliprotein Content in Cyanobacterium Synechocystis. J. Vis. Exp. 1–9.

Zavřel, T., Faizi, M., Loureiro, C., Sinetova, M., Zorina, A., Poschmann, G., Stühler, K., Steuer, R., Červený, J., 2019. Quantitative insights into the cyanobacterial cell economy. Elife 8, 1–29.

Zavřel, T., Očenášová, P., Červený, J., 2017. Phenotypic characterization of Synechocystis sp. PCC 6803 substrains reveals differences in sensitivity to abiotic stress. PLoS One 12, e0189130.

Zavřel, T., Očenášová, P., Sinetova, M., Červený, J., 2018b. Determination of Storage (Starch/Glycogen) and Total Saccharides Content in Algae and Cyanobacteria by a Phenol-Sulfuric Acid Method. Bio-Protocol 8, 1–13.

Zavřel, T., Schoffman, H., Lukeš, M., Fedorko, J., Keren, N., Červený, J., 2021. Monitoring fitness and productivity in cyanobacteria batch cultures. Algal Res. 56, 1–15.

Zavřel, T., Sinetova, M.A., Búzová, D., Literáková, P., Červený, J., 2015a. Characterization of a model cyanobacterium Synechocystis sp. PCC 6803 autotrophic growth in a flat-panel photobioreactor. Eng. Life Sci. 15, 122–132.

Zavřel, T., Sinetova, M.A., Červený, J., 2015b. Measurement of Chlorophyll a and Carotenoids Concentration in Cyanobacteria. bio-protocol 5, 1–5.

Zavřel, T., Szabó, M., Tamburic, B., Evenhuis, C., Kuzhiumparambil, U., Literáková, P., Larkum, A.W.D., Raven, J.A., Červený, J., Ralph, P.J., 2018c. Effect of carbon limitation on photosynthetic electron transport in Nannochloropsis oculata. J. Photochem. Photobiol. B Biol. 181.

