## Supplementary Material for "A comprehensive study of light quality acclimation in *Synechocystis* sp. PCC 6803"

**for**

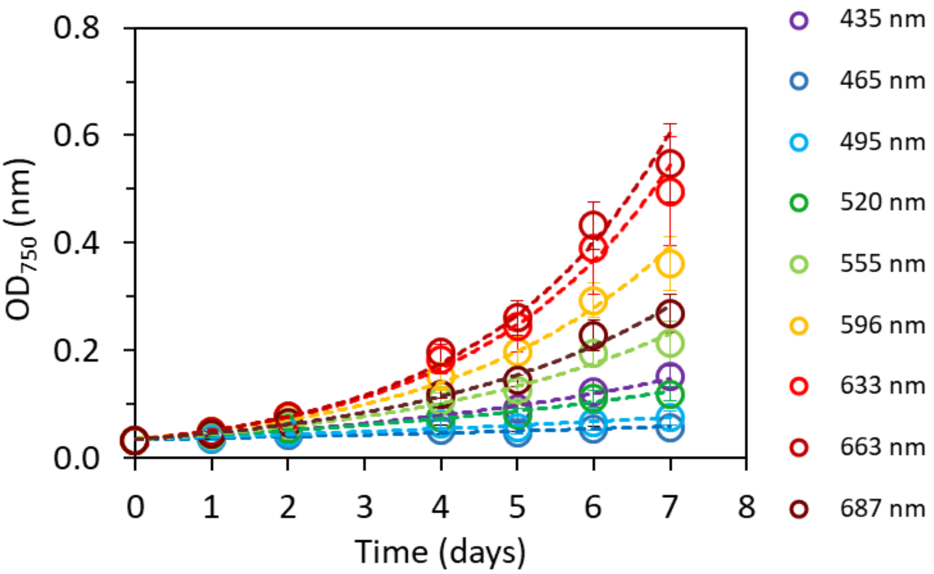


**Figure S1.** Measurements of optical density at 750 nm ($OD_{750}$) from which the specific growth rates were estimated. The values represent mean±SD, n=4. The dashed lines represent the data extrapolation by exponential regression model ($R^2$ varied between 0.89–1.00).

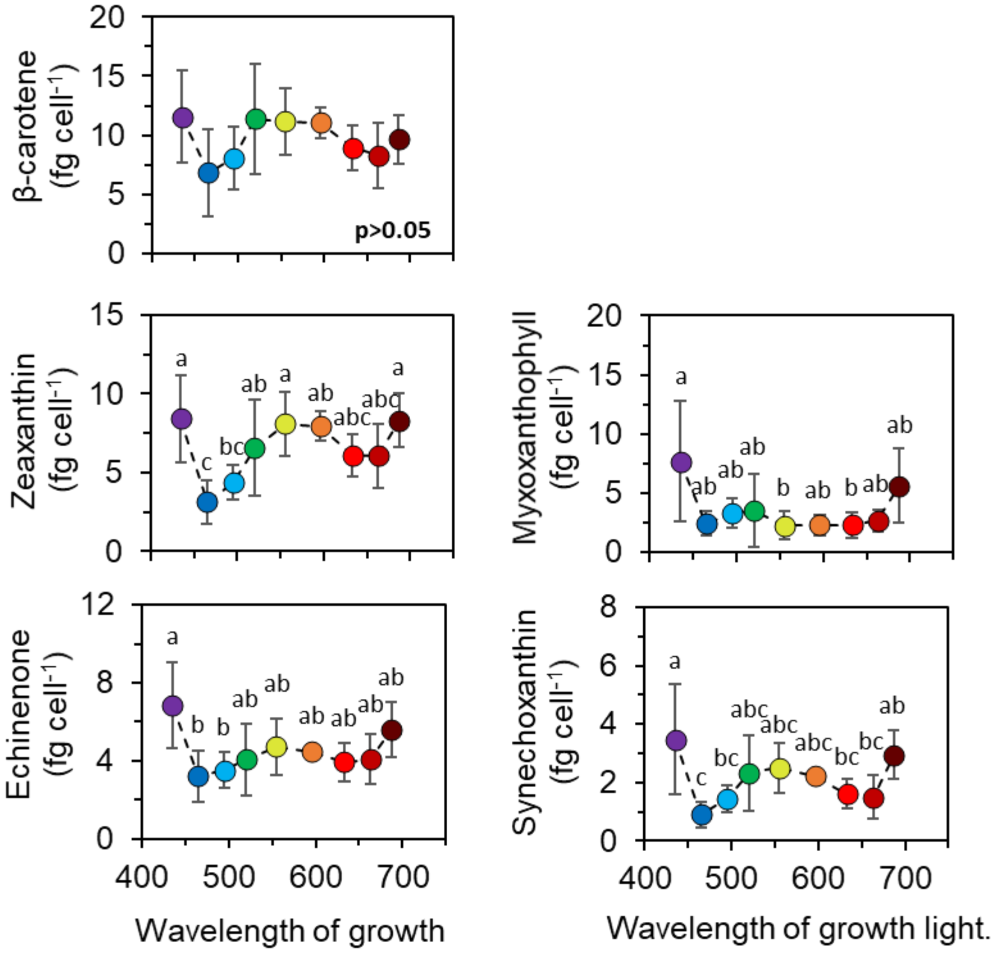


**Figure S2.** Abundance of the five major carotenoids in *Synechocystis* cells. The values represent mean±SD (n=3). The letters above the symbols indicate statistically significant differences within each parameter ($p<0.05$).

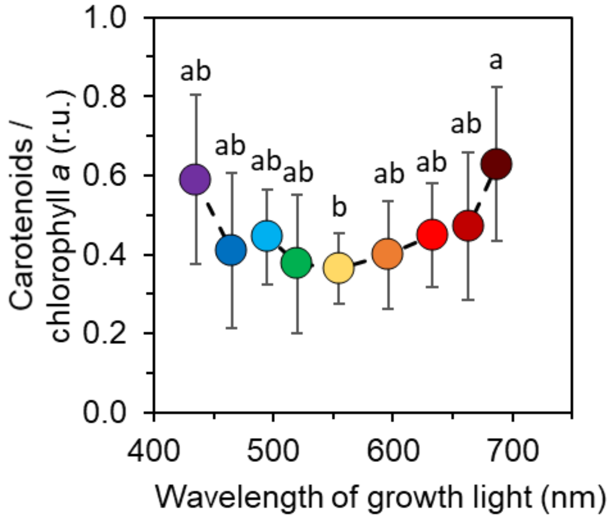


**Figure S3.** Mass ratio of total carotenoids and chlorophyll *a* in *Synechocystis* cells. The values represent mean±SD (n= 3). The letters above the symbols indicate statistically significant differences within each parameter ($p<0.05$).

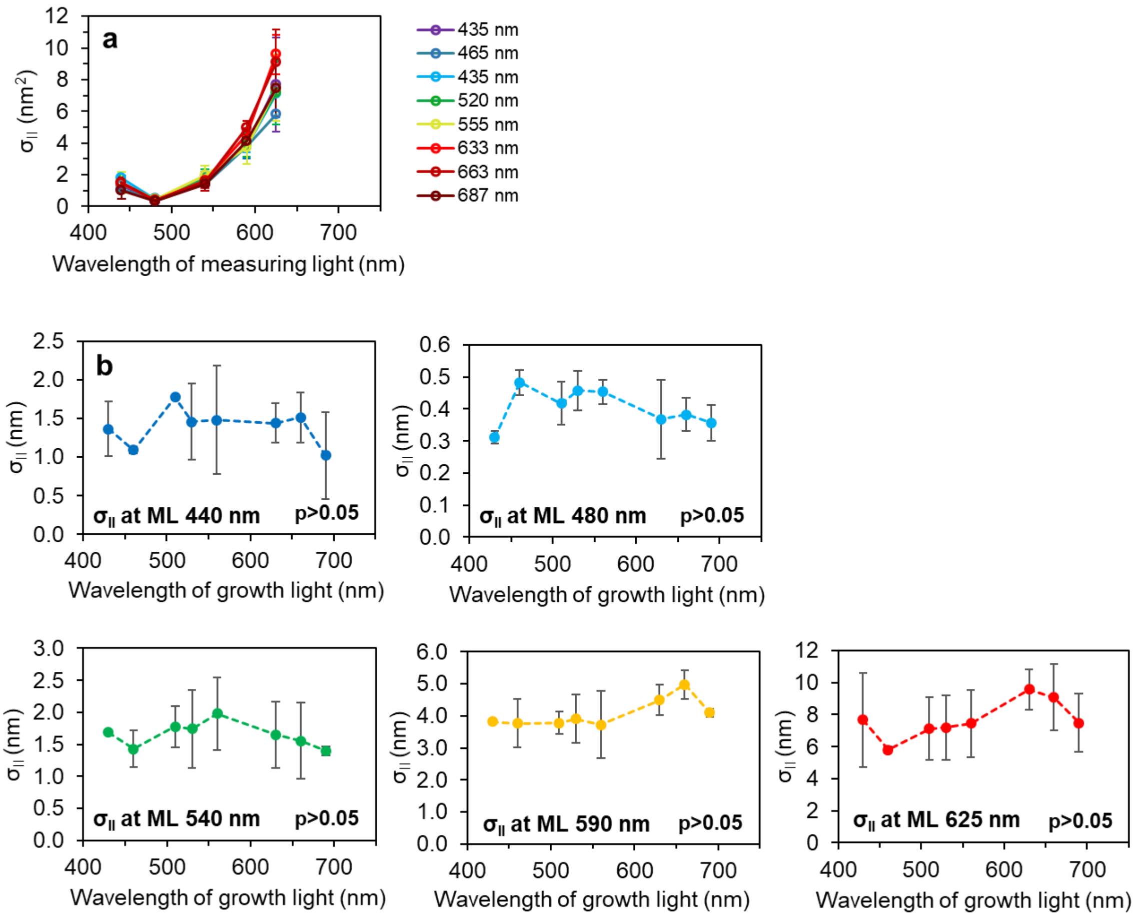


**Figure S4.** Functional absorption cross-section of PSII ($\sigma_{II}$) in *Synechocystis* cells. **a:** $\sigma_{II}$ as a function of wavelength of measuring light (ML) set by Multi-Color-PAM [1]. **b:** $\sigma_{II}$ as a function of wavelength of cultivation light. All values represent mean±SD (n=3). The p-values indicate the result of the Kruskal-Wallis test.

a

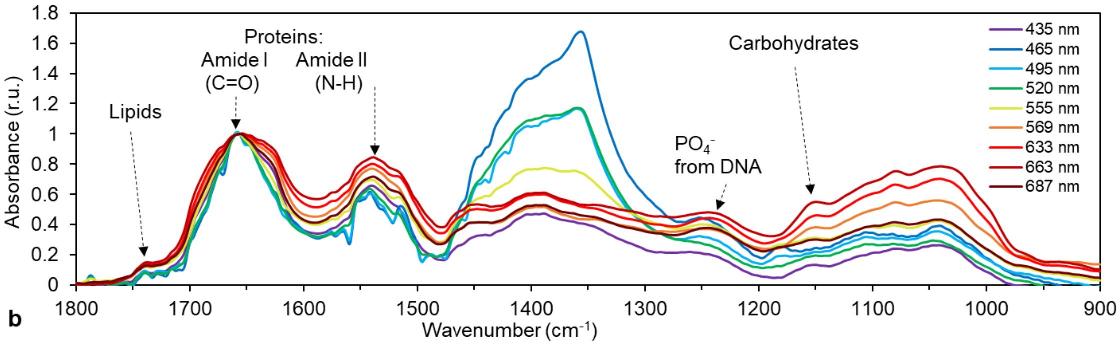


**Figure S5.** FTIR spectra of *Synechocystis* cultures, normalized to Amide I peak (1652 cm$^{-1}$). The spectra represent the mean of three biological replicates; error bars are omitted for clarity.

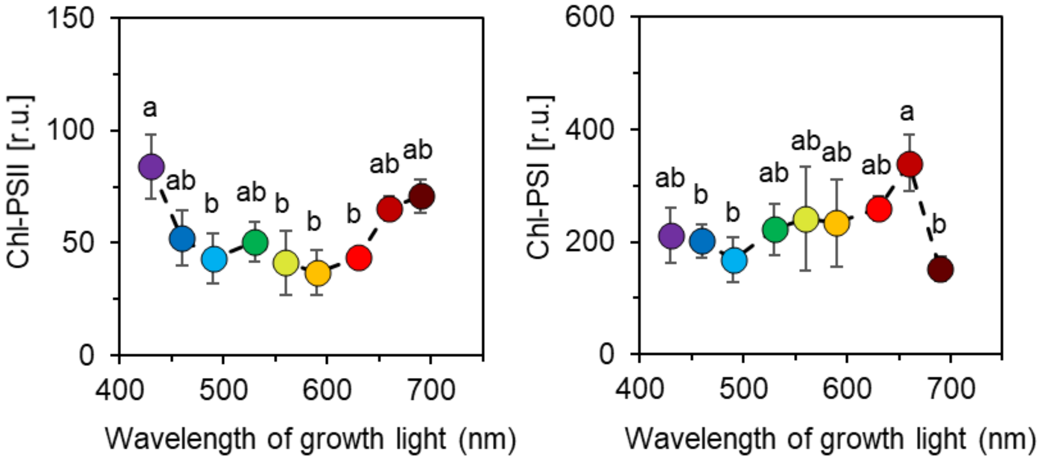


**Figure S6.** Estimation of PSII and PSI fluorescence in *Synechocystis* cells, based on low temperature (77K) Chl *a* fluorescence emission at 695 nm and 726 nm, respectively, upon excitation at 440 nm and corrected for fluorescence emission from PBS-PSII and PBS-PSI (Eq. 9-10). The values represent mean±SD (n=3). The letters above the symbols indicate statistically significant differences within each parameter ($p<0.05$).

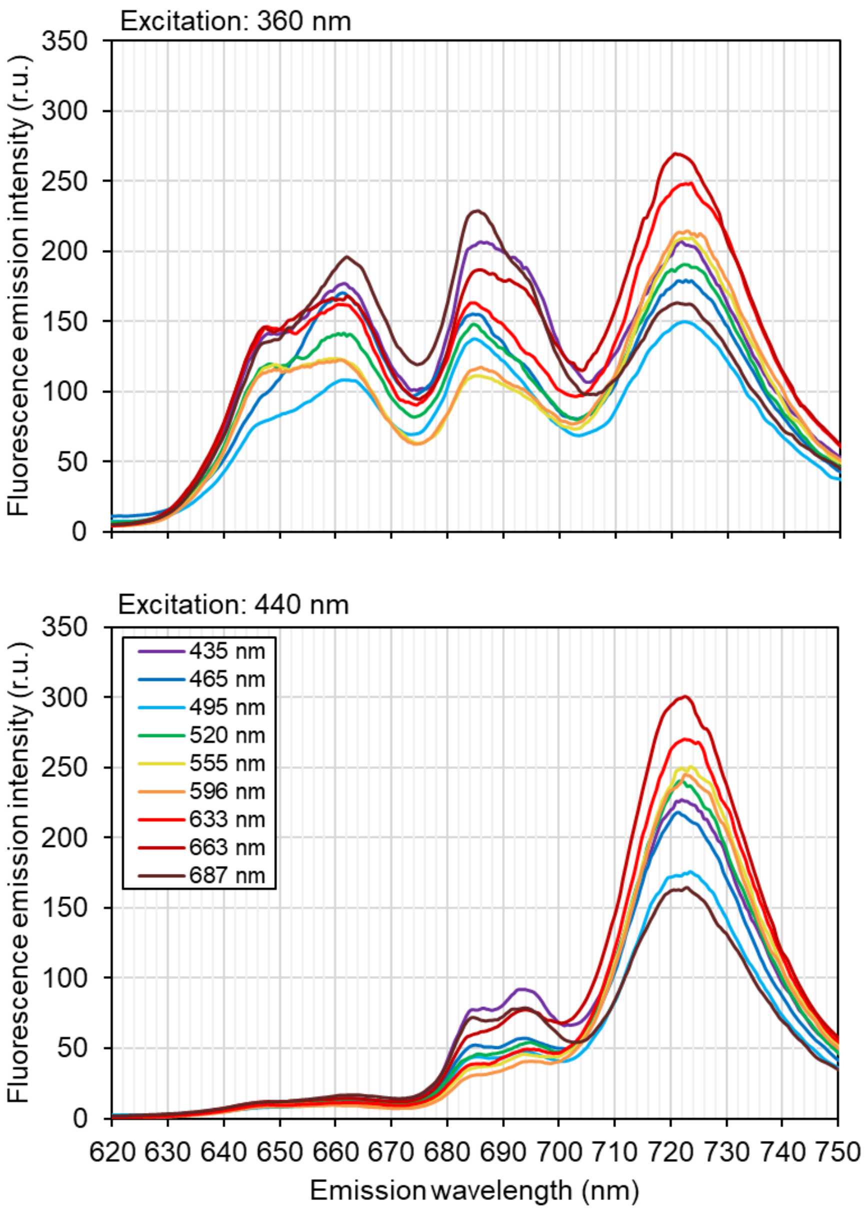


**Figure S7.** Fluorescence emission spectra of *Synechocystis* cultures recorded upon 360 nm (upper panel) and 440 nm excitation (bottom panel) at 77K. The spectra represent the mean of three biological replicates; error bars are not shown for clarity.

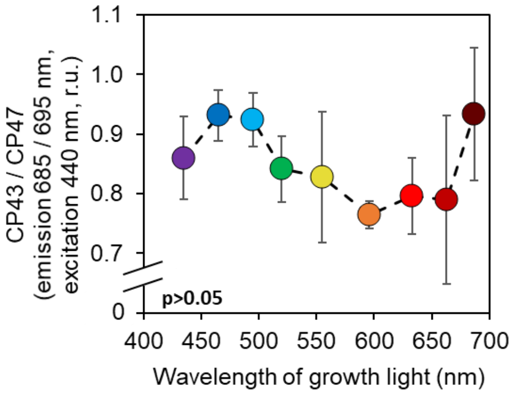


**Figure S8.** Estimation of the 685 / 695 nm fluorescence ratio in *Synechocystis* cells upon 440 nm excitation, as a proxy for the ration of CP43 / CP47. The fluorescence values were corrected for emission from PBS-PSII (Eq. 9-10). The values represent mean±SD (n=3). The p-values indicate result of Kruskal-Wallis test.

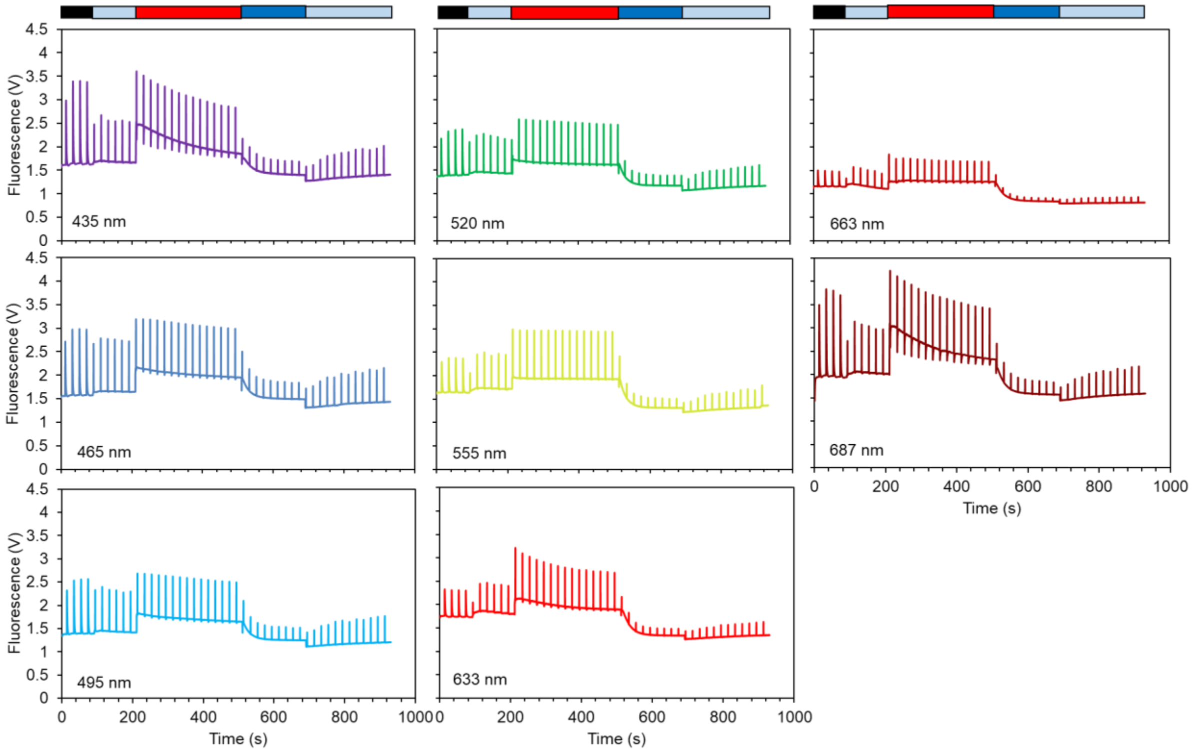


**Figure S9.** Fluorescence induction curves recorded to estimate LEF, the rate of *State 1* → *State 2* transition, and NPQ in *State 2* in *Synechocystis* cultures. The color bars above the charts represent the dark acclimation period (black) and four actinic light periods of different wavelengths and intensities: 480 nm AL with an intensity of either 80 µmol photons m$^{-2}$ s$^{-1}$ (light blue, 2x) or 1 800 µmol photons m$^{-2}$ s$^{-1}$ (dark blue), and 625 nm AL with an intensity of 50 µmol photons m$^{-2}$ s$^{-1}$ (red). The spectra represent the mean of three biological replicates; error bars are omitted for clarity.

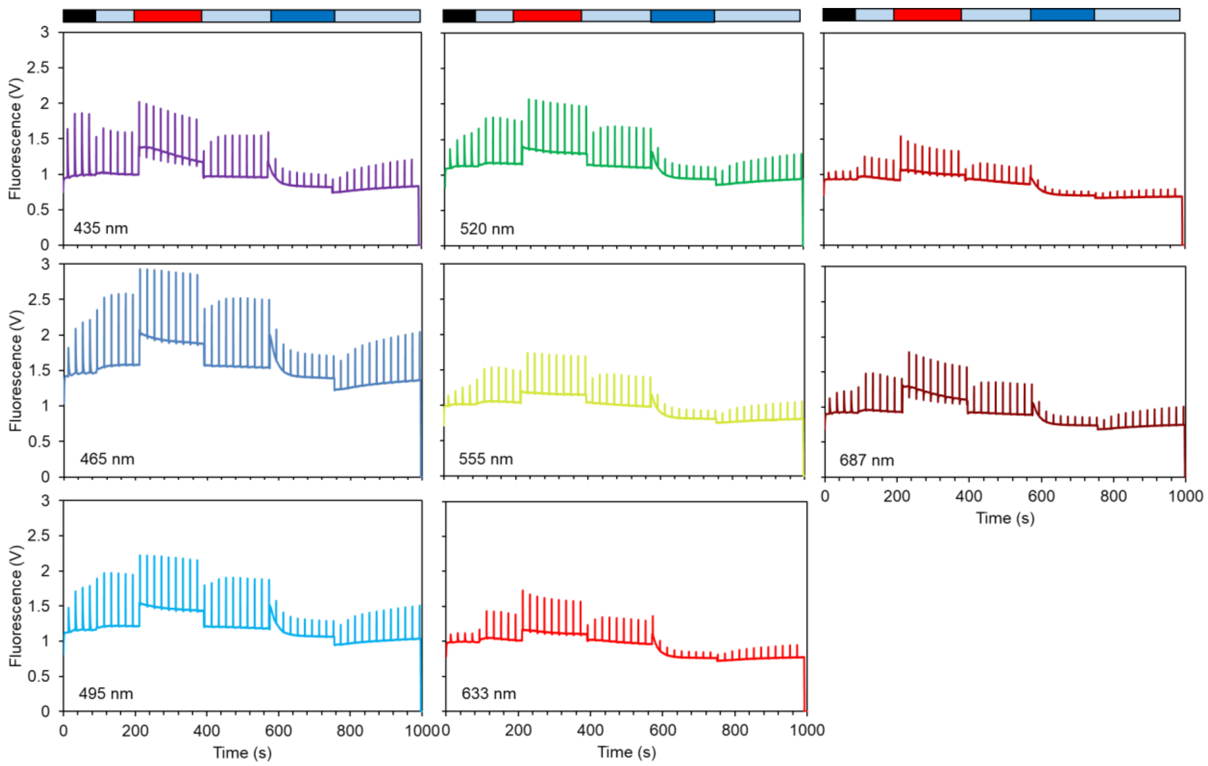


**Figure S10.** Fluorescence induction curves recorded to estimate LEF, rate of *State 2* → *State 1* transition, and NPQ in *State 1* in *Synechocystis* cultures. The color bars above the charts represent the dark acclimation period (black) and five actinic light periods of different wavelengths and intensities: 480 nm AL with an intensity of either 80 µmol photons $m^{-2}$ $s^{-1}$ (light blue, 3x) or 1 800 µmol photons $m^{-2}$ $s^{-1}$ (dark blue), and 625 nm AL with an intensity of 50 µmol photons $m^{-2}$ $s^{-1}$ (red). The spectra represent the mean of three biological replicates; error bars are omitted for clarity.

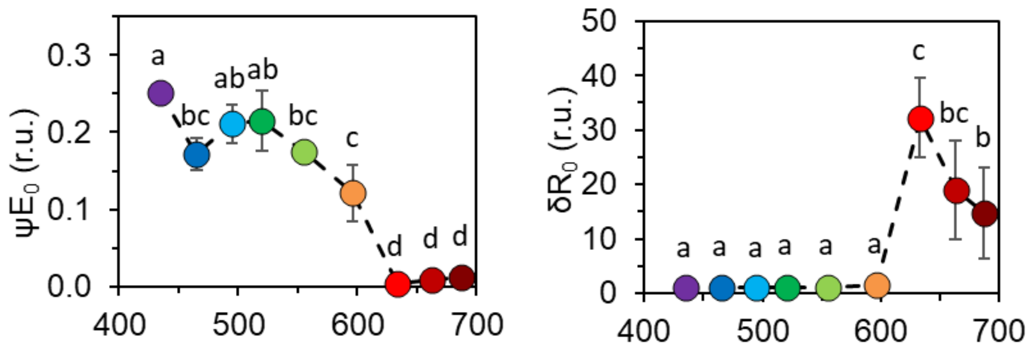


**Figure S11.** Parameters ψ$E_0$ and δ$R_0$, as derived from OJIP curves (Figure 5) in light-acclimated *Synechocystis* cultures, representing the efficiency with which a PSII trapped electron was transferred from $Q_A^-$ to PQ (ψ$E_0$, left) and the efficiency with which an electron from $PQH_2$ was transferred to final PSI acceptors (δ$R_0$, right) [2]. The values represent mean±SD (n=3). The letters above the symbols indicate statistically significant differences within each parameter ($p<0.05$).

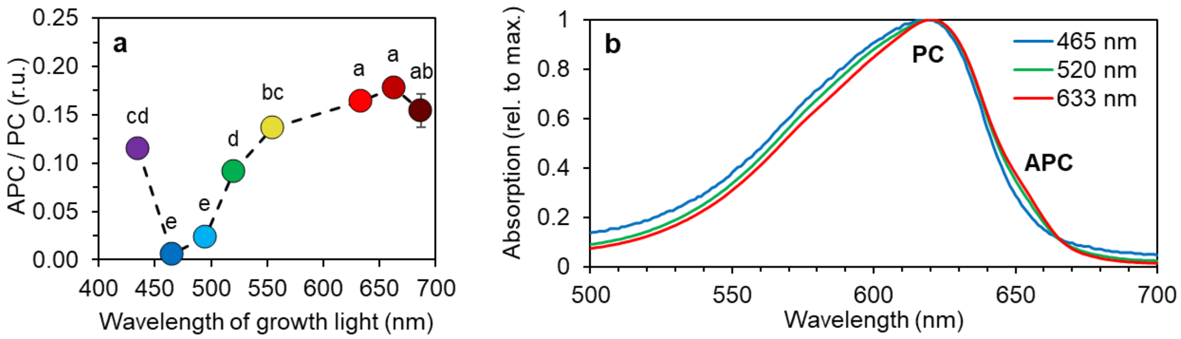


**Figure S12. a:** Allophycocyanin / phycocyanin ratio in *Synechocystis* cells. The values represent the mean±SD (n=3). The letters above the symbols indicate statistically significant differences within each parameter ($p<0.05$). **b:** Absorption spectra of PBS extracts [3] of cultures cultivated under 465 nm, 520 nm and 633 nm lights, as the mean of three biological replicates; error bars are omitted for clarity.

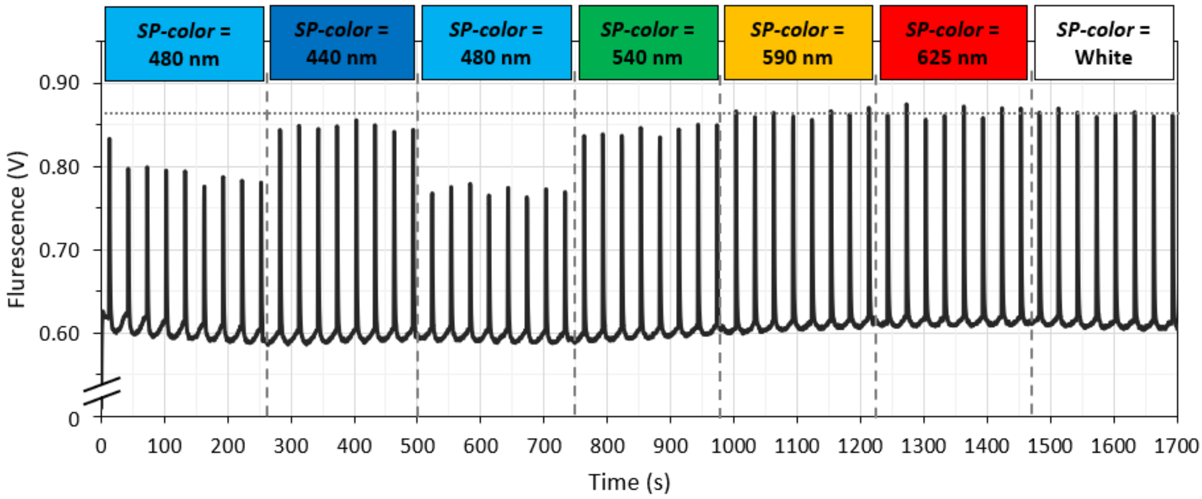


**Figure S13** A chlorophyll fluorescence trace of *Synechocystis* upon consecutively using saturation pulses (SP-color) with different colors, set by changing the *AL-color* setting at the user-defined protocol of MULTI-COLOR-PAM. During the measurement, Actinic Light (AL) was turned off, and Measuring Light (ML), Damping and Gain were set to keep $F_0$ around the value of 0.6. The ML color and the length of the SP (*SP-width*) was set to 625 nm and 600 ms, respectively. The only varied parameter, *AL-color*, defined the color of the applied SP. The SP intensity was set to the maximum for each wavelength (*SP-int=20*). Prior to the measurement, the *Synechocystis* culture was cultivated in BG-11 medium in Erlenmeyer flask on a shaker at 23 °C under cool white light of 30 µmol photons $m^{-2}$ $s^{-1}$ on air. Culture density during the measurement was $OD_{750}$ = 0.2. Prior to the measurement, 1.5 mL culture was transferred to a quartz cuvette and dark-acclimated for 10 min. Dashed lines indicate shifts in *SP-color* applied in the Multi-Color PAM protocol. Th dotted line indicates maximal fluorescence level reached under 590 nm, 625 nm and white AL.

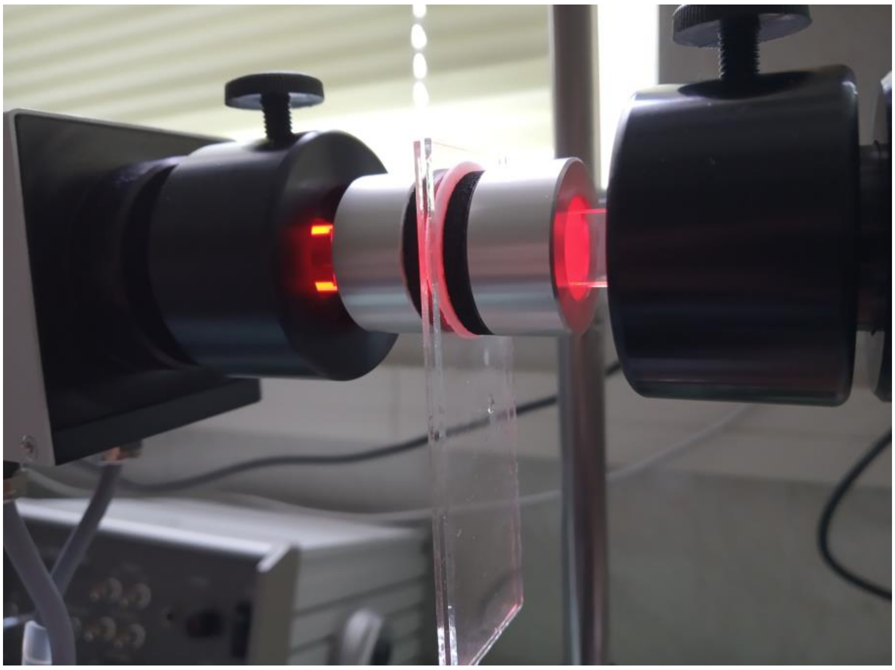

**Figure S14.** Experimental setup for P700 kinetic measurements by Dual-PAM-100. A wet GF/B filter with cells was placed between two microscopy glass slides within the leaf holder DUAL-B.

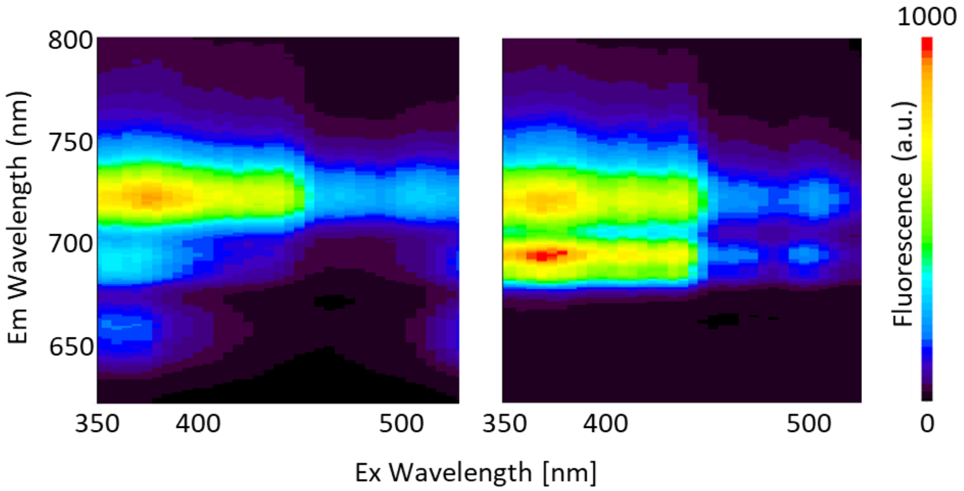


**Figure S15.** Representative excitation-emission fluorescence maps recorded at 77K for *Synechocystis* wild-type (WT) (left) and PBS-less PAL mutant [4] (right).

**References:**

1. Schreiber U, Klughammer C, Kolbowski J. Assessment of wavelength-dependent parameters of photosynthetic electron transport with a new type of multi-color PAM chlorophyll fluorometer. *Photosynth Res* 2012; **113**: 127–144.

2. Stirbet A, Lazár D, Kromdijk J, Govindjee. Chlorophyll a fluorescence induction: Can just a one-second measurement be used to quantify abiotic stress responses? *Photosynthetica* 2018; **56**: 86–104.

3. Zavřel T, Chmelík D, Sinetova MA, Červený J. Spectrophotometric Determination of Phycobiliprotein Content in Cyanobacterium Synechocystis. *J Vis Exp* 2018; 1–9.

4. Ajlani G, Vernotte C. Construction and characterization of a phycobiliprotein-less mutant of Synechocystis sp. PCC 6803. *Plant Mol Biol* 1998; **37**: 577–580.
